# Diverse bacteria exhibit superadditive growth on mixed classes of carbon sources

**DOI:** 10.64898/2026.11.06.764671

**Authors:** Elena K. Perry, Ying Wang, Valentine V. Trotter, Yan Chen, Markus de Raad, Suzanne M. Kosina, Rowan Nelson, Christopher J. Petzold, Daniel Segrè, Adam P. Arkin, Adam M. Deutschbauer, Trent R. Northen

## Abstract

Heterotrophic bacteria in nature encounter complex mixtures of functionally-distinct carbon sources, yet whether such diversity enhances growth yield across diverse taxa remains unknown. We grew dozens of strains from four phyla on defined mixtures representing five classes of carbon sources. Remarkably, many but not all strains exhibited superadditive growth on a dilute mixture of all 64 compounds—that is, yield exceeding the weighted sum of yields on more concentrated subsets of functionally-similar compounds. Using exometabolomics, pooled genetic screens, and proteomics, we find that superadditive growth is associated with enhanced carbon consumption in the mixed-class medium, combined with reduced dependence on amino acid and nucleotide biosynthesis. Moreover, superadditive growth was predictable from genomes within the tested strain panel and correlated with genomically encoded sugar-over-acid preference. These findings reveal a previously underrecognized axis of physiological variation among heterotrophs and open the door to understanding the biochemical and evolutionary basis of this divergence.

---

Ever since Monod’s seminal experiments in the 1940s, it has been accepted that under carbon-limited conditions, the batch growth yield of a heterotrophic bacterium on a single carbon source will generally be proportional to the initial compound concentration^1,2^. Moreover, if two similar carbon sources are mixed, the resulting biomass yield is typically the sum of the yields produced on each^1^. In nature, however, heterotrophic bacteria frequently encounter a vast array of organic compounds belonging to different functional classes (e.g. sugars, amino acids, organic acids, etc.), though they are often still carbon-limited^3,4^. To what extent is growth yield an additive property under such conditions?

Despite decades of bacterial growth studies, the answer to this important question remains poorly understood. Growth yield, as opposed to growth rate, is likely a key driver of evolutionary success in fragmented, resource-limited habitats such as soil^5,6^; growth yield may also impact community dynamics in host-associated environments, such as the gut mucosa^7^; and microbial biomass production contributes significantly to long-term stabilization of organic carbon both in soil and in the ocean^8,9^. Furthermore, biomass yield on complex feedstocks is directly relevant to the economics of biomanufacturing^10,11^. Thus, an improved understanding of growth yield on mixed substrates could impact multiple subfields of microbiology.

Previous studies have reported several forms of nonadditive response to substrate mixtures. In a natural freshwater bacterial community, mixtures of dissolved organic carbon sources increased bacterial growth efficiency, measured from biomass production and respiration, relative to expectations derived from individual sources^12^. In a marine model system, *Ruegeria pomeroyi* attained higher optical density on mixed substrates than predicted from the weighted sum of single-substrate cultures under a daily growth–dilution regime^13^. Experiments in *Escherichia coli* further showed that the contribution of a nutrient to batch biomass production can depend on the identity and initial amount of another nutrient^2^, and a similar cooperativity in promoting growth was also predicted for pairs of carbon sources based on genome-scale models^14^. These findings motivate testing how commonly diverse bacteria depart from an explicit additive biomass expectation and exploring how mixed substrate growth may contribute to higher growth yields.

Here, we propose the term “superadditive” growth to describe cases where biomass yield on a mixture of substrates exceeds the weighted sum of yields from subsets of the full mixture. Using a combination of high-throughput growth characterization, multi-omics, and machine learning, we set out to address three questions: 1) How common is superadditive growth on dilute mixtures of functionally-distinct carbon sources across diverse heterotrophic bacteria? 2) What mechanisms most likely underpin superadditive growth? 3) Can the impact of mixing different classes of carbon sources on biomass yield be predicted from genomic traits?

## Results

### Diverse bacteria achieve superadditive growth yields on dilute mixed classes of substrates

We first assembled a collection of 98 bacterial strains isolated from diverse soils and sediments, along with two *E. coli* strains as representative model organisms that have been extensively studied (Supplementary Table 1). We grew each strain individually in six defined media containing different combinations of 64 carbon sources commonly found in soils (Fig. 1a, Supplementary Table 2)^15^. We divided these carbon sources into five classes: sugars, proteinogenic amino acids, organic acids, nucleobases/sides, and “other” (compounds that do not fit neatly into one of the other categories). The single-class media, Sugar7, AminoAcid20, OrganicAcid7, Nucleo14, and Other16, each contained the class and number of carbon sources indicated in the name, while All64 contained all of the carbon sources. In each medium, the carbon sources were added at equimolar concentrations while keeping the total concentration constant at 2.56 mM (see Methods for rationale).

**Figure 1.**
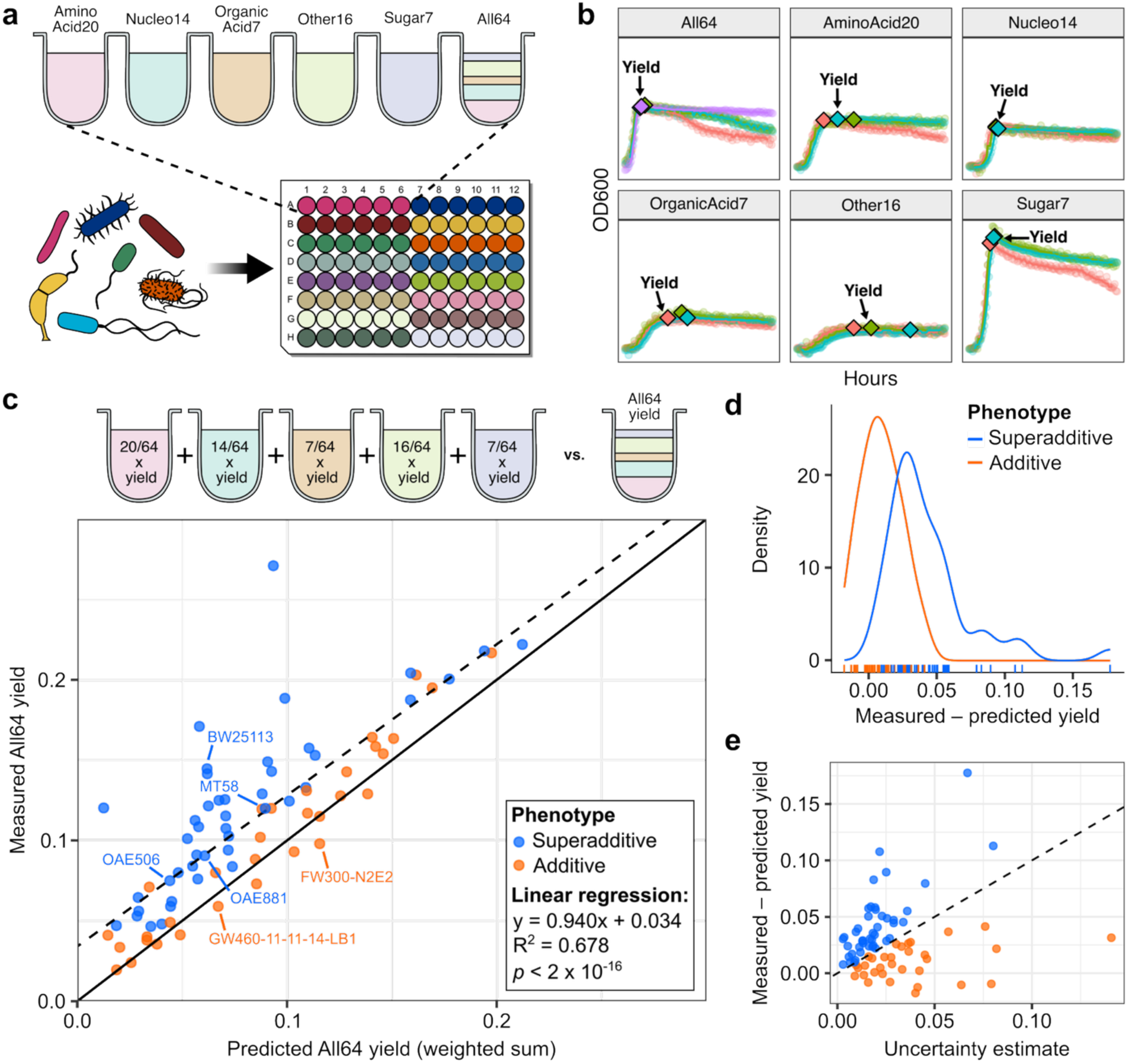
Diverse bacteria achieve superadditive growth yields on mixed classes of carbon sources. **a)** Experimental setup for high-throughput within-strain comparison of growth on single-class versus mixed-class carbon sources for diverse heterotrophic bacteria. Each single-class medium contained the class and number of carbon sources in the name while All64 contained all of the carbon sources from the five single-class media. In each medium, the carbon sources were provided at equimolar concentrations summing to 2560 µM. Strains were grown individually in 96-well plate format for 64 hrs, with 2-5 replicates per strain and condition. **b)** Example determination of yield from growth curves. The colors represent independent replicates of the same strain performed on separate days. The median of the maximum OD_600_ from the smoothed growth curve (solid line overlaid on the individual data points) was taken as the representative yield for each combination of strain and medium. **c)** Plot of measured growth yield on All64 versus the weighted-sum prediction. The cartoon depicts the comparison being made. Each data point represents the median value for an individual strain. The solid line is y = x (the baseline expectation) and the dashed line is the best-fit linear regression model whose equation is given on the bottom right. Labeled strains were used in follow-up experiments. **d)** Probability density function for the difference between measured and predicted yield on All64, split by the assigned phenotype. Tick marks represent the individual underlying data points. The difference is greater on average for superadditive strains as expected, but the distributions for the two phenotypes partially overlap because our definition of superadditivity also accounts for the variability between replicates for each strain, as shown in panel (e). **e)** Plot of the difference between measured and predicted yield on All64, versus the total uncertainty estimate for each strain (sum of the scaled median absolute deviation for All64 and the propagated error estimate for the weighted sum prediction; see Materials and Methods for details), enabling visualization of our cutoff for defining a strain as a “superadditive” grower: the difference between measured and predicted yield on All64 must exceed the uncertainty estimate for that strain. Data points are colored by the assigned phenotype. The dashed line is y = x.

The strains were grown in 96-well plates and optical density at 600 nm (OD_600_) was tracked for 64 hours, by which time most strains had reached stationary phase. Growth yield was defined as the maximum OD_600_ value of each growth curve after smoothing to mitigate noise (Fig. 1b; see Methods). Among the 100 strains tested, 24 strains failed to grow either in pre-cultures or in any of the experimental media and were excluded from subsequent analyses. For the remaining 76 strains, which represent 30 taxonomic families, comparing the measured yield on All64 versus the weighted sum of yields on the five single-class media revealed that the majority of strains achieved higher-than-additive (“superadditive”) yields on All64 (Fig. 1c), with a median relative yield gain of 31% over the additive expectation, even though All64 was not the highest-yielding medium in absolute terms for most strains (Extended Data Fig. 1).

To facilitate downstream analyses, we developed a pipeline for categorizing the strains into putative “superadditive” versus “additive” growth phenotypes. Rather than setting a fixed threshold, we sought to identify strains whose median yield on All64 substantially exceeded the weighted sum prediction after accounting for the variability among biological replicates (see Methods). Forty-four strains (58%) met our criteria for superadditive growth; the rest were classified as additive (Fig. 1c, Supplementary Table 3). While the distributions of the superadditive and additive growers partially overlap in terms of difference between observed and expected median yield on All64 (Fig. 1c-d), this is because our criteria for superadditivity required strains with higher variability among replicates to exceed the additive expectation by a greater difference (Fig. 1e, Supplementary Table 3).

To assess whether superadditive versus additive growth is related to phylogeny, we constructed a phylogenomic tree for the 72 strains for which genome sequences were available. We found that these phenotypes are largely consistent within various clades. However, the taxonomic level at which it is conserved appears to vary (Extended Data Fig. 2), and at least seven pairs of closely-related strains within the same genus or species exhibited discordant phenotypes (Extended Data Fig. 3).

We next asked which physiological differences accompany departures from the concentration-scaled additive growth prediction. Two potentially contributing processes are increased utilization of supplied substrates and reduced investment in biosynthesis when biomass building blocks are externally available. To explore these possibilities, we selected six strains for follow-up investigations based on the availability of genome-wide transposon mutant libraries. In particular, we used exometabolomics to assess net depletion of supplied compounds, pooled genetic assays to examine conditional biosynthetic dependencies, and proteomics to assess biosynthetic enzyme expression. Four of the strains (*E. coli* BW25113, *Pantoea* sp. MT58, *Lysobacter* sp. OAE881, and *Bosea* sp. OAE506) represent the superadditive growth phenotype, while the remaining two strains (*Hydrogenophaga* sp. GW460-11-11-14-LB1 and *Pseudomonas fluorescens* FW300-N2E2) represent the additive growth phenotype (Fig. 1c).

### Superadditive growers consumed considerably more carbon than expected in the mixed-class medium

First, to address whether mixing different classes of carbon sources stimulates additional substrate consumption compared with single-class media, we performed exometabolomic profiling^16^. The representative strains were grown to early stationary phase in all six media and the provided carbon sources were quantified in the spent supernatants relative to uninoculated controls. Our null hypothesis was that in carbon-limited media such as those used in this study, all carbon sources that a strain can utilize would be essentially fully consumed as of entry into stationary phase. Thus, while All64 starts with a lower concentration of each substrate, the percent remaining for each usable substrate after growth would theoretically be similar (∼0%) to the single-class media in the absence of co-metabolism-like effects.

In contrast to this baseline expectation, we found that all of the representative superadditive growers depleted numerous compounds (16-34 per strain) to a significantly greater proportional extent in All64 relative to the respective single-class media (Fig. 2, Supplementary Table 4), of which the majority were amino acids or nucleobases/sides. On the other hand, the representative additive growers each consumed only 8-11 compounds proportionally more in All64, of which just 2-5 were amino acids or nucleobases/sides. These trends were similar for compounds that were consumed more in absolute terms in All64 (Extended Data Fig. 4). Even though different substrates yield different amounts of energy per mole and require different levels of proteomic investment to metabolize, total net depletion of the supplied carbon sources correlated well with growth yield (Extended Data Fig. 5) (Supplementary Notes). Indeed, while the total net carbon source depletion in All64 (see Methods) exceeded the weighted sum prediction to some degree for all six strains, this discrepancy was considerably larger for the superadditive growers (Fig. 2b). Taken together, the exometabolomics results are consistent with increased substrate utilization contributing to the superadditive growth response.

**Figure 2.**
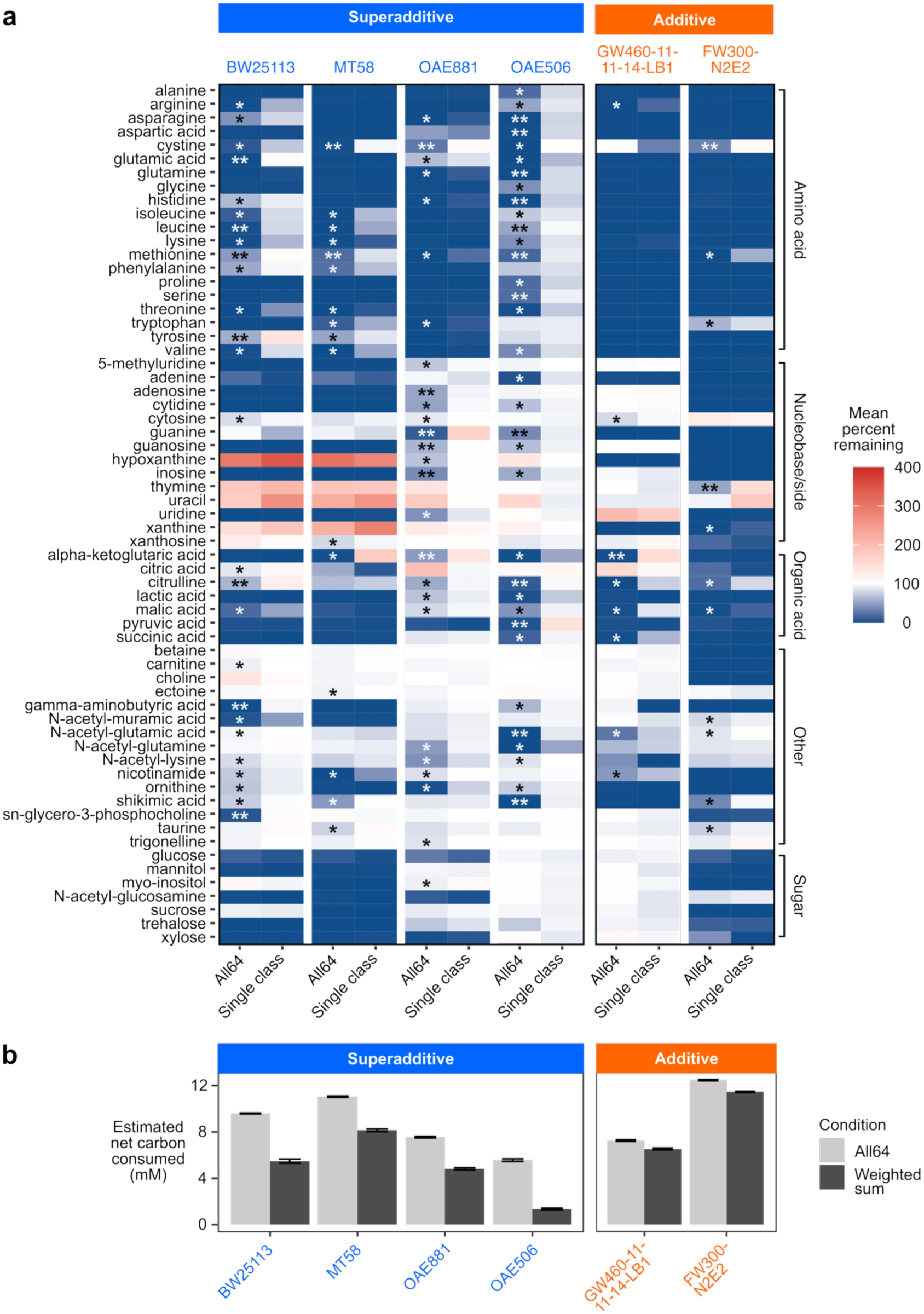
Superadditive growers consumed much more carbon than expected in the mixed-class medium. **a)** Heat map depicting the mean percent remaining of 63 carbon sources (spermidine was not detectable) in the mixed metabolite class medium (All64) versus the respective single-class medium (AminoAcid20, Nucleo14, OrganicAcid7, Other16, or Sugar7) in the supernatants of cultures grown to early stationary phase. 100% remaining (white) indicates no net consumption; <100% (blue) indicates net consumption; and >100% (red) indicates net excretion. *Proportional consumption was significantly greater in All64 than the single-class medium (*p* < 0.05, Welch’s t-test followed by the Benjamini-Hochberg procedure, excluding compounds with net excretion in All64 or less than 5% difference between All64 versus the single-class medium); **net absolute consumption was also significantly greater in All64 (*p* < 0.05, Welch’s t-test followed by the Benjamini-Hochberg procedure). For representation of the net absolute amount consumed for each compound, see Extended Data Fig. 4. **b)** Mean estimated total net carbon consumption in All64 in the exometabolomics experiment, based on depletion of provided carbon sources, compared to the weighted sum of estimated net carbon consumption in the five single-class media. Error bars represent the standard deviation (n = 3). For all strains, the measured estimated net carbon consumption was significantly higher than the weighted sum prediction (*p* < 0.001, Welch’s t-test followed by the Benjamini-Hochberg procedure).

### High-throughput genetic screens reveal differences in the importance of biosynthetic pathways

To identify potential genetic mechanisms underpinning superadditive versus additive growth, we next turned to high-throughput genetic screening via random barcode transposon-site sequencing (RB-TnSeq), which allows thousands of uniquely barcoded transposon insertion mutants to be pooled and grown competitively under numerous experimental conditions^17^. With this method, negative fitness scores indicate that a gene is important for growth under the experimental condition, while positive fitness scores indicate that disruption of the gene improves relative fitness in the pooled assay. We obtained RB-TnSeq libraries for all six representative strains; four have previously been published^17–20^, while the libraries for *Bosea* OAE506 and *Hydrogenophaga* GW460-11-11-14-LB1 are reported here for the first time. After recovering the libraries in rich media, we performed experiments in triplicate in All64 and as many single-class media as supported robust growth. We did not find putative regulatory genes with negative fitness scores specific to All64 for any of the strains. However, a few notable patterns of fitness effects emerged.

First, the superadditive growers consistently exhibited fewer dependencies on amino acid and nucleotide biosynthesis genes in All64 compared with the additive growers (Fig. 3, Supplementary Table 5, Supplementary Table 6). This indicates that the low concentrations of amino acids and nucleobases/sides in All64 medium rescued a greater variety of auxotrophic mutants for superadditive relative to additive strains. Second, among the four strains with high-quality data from at least five of the six media, we found that the two additive growers each had a gene (pyruvate dehydrogenase E1 component in *Hydrogenophaga* GW460-11-11-14-LB1, and a maltose/maltodextrin transporter in *P. fluorescens* FW300-N2E2) whose disruption significantly improved fitness specifically in All64, whereas the superadditive growers had no such genes (Supplementary Table 5). Finally, in all strains, multiple catabolic genes had significant negative fitness scores in All64 (Supplementary Table 5), but no consistent patterns were observed between superadditive and additive strains regarding the fitness effects of particular catabolic pathways (Supplementary Notes).

**Figure 3.**
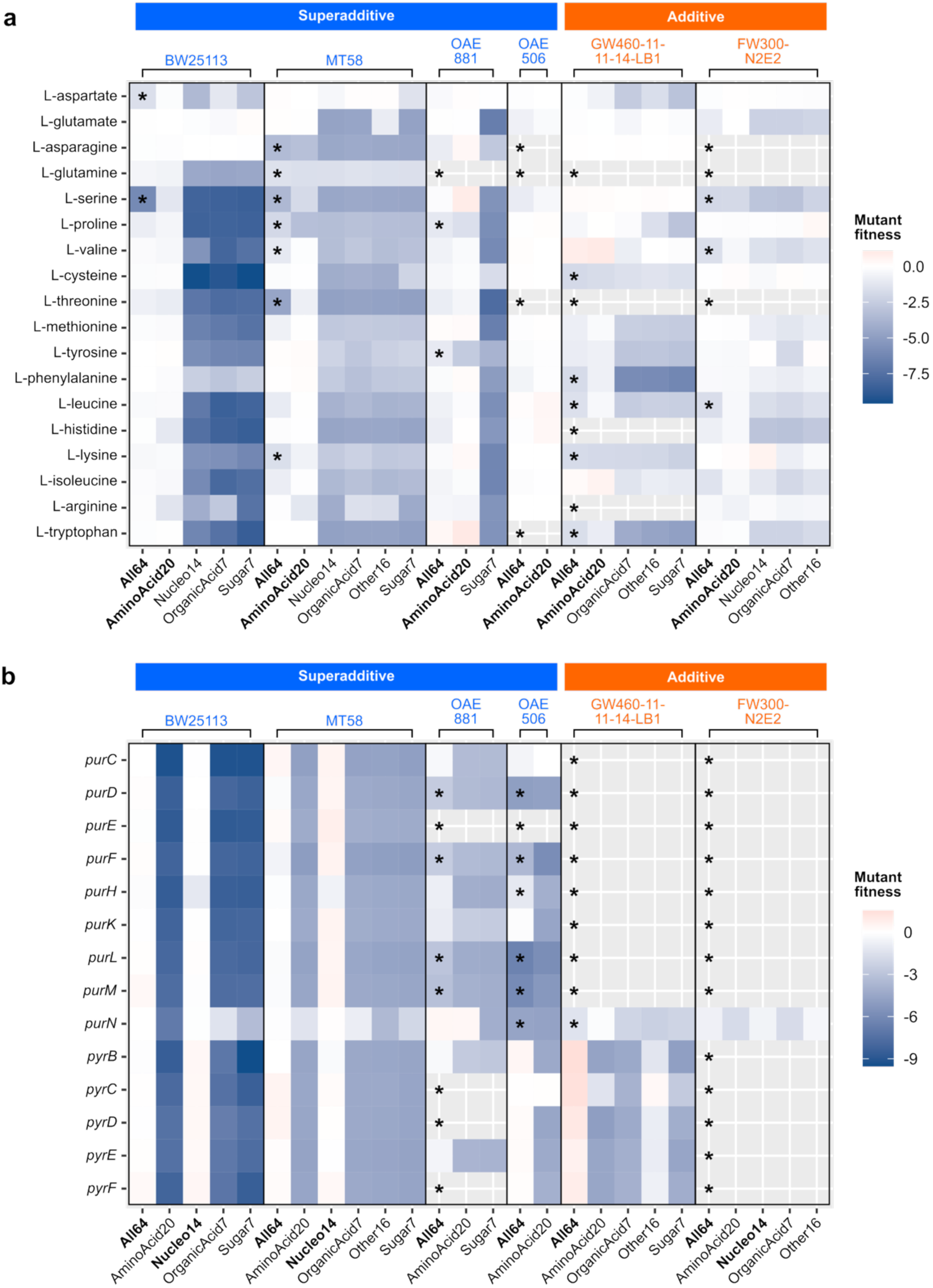
RB-TnSeq reveals differences in the fitness effects of biosynthetic pathways. **a)** Heat maps of mean mutant fitness score per amino acid biosynthetic pathway across the different media, separated by strain and annotated by growth phenotype on All64 (superadditive vs. additive). Alanine and glycine were excluded due to ambiguity in pathway analysis (see Methods). Media containing amino acids are labeled in bold font. Asterisks mark pathways whose constituent genes were either essential based on TnSeq analysis or did not have detectable mutants at Time0 after recovery in rich media (blank rows), or had a mean mutant fitness score < –1 in All64. **b)** Heat maps of mutant fitness scores for *de novo* nucleotide biosynthesis genes across the different media, separated by strain and annotated by growth phenotype on All64 (superadditive vs. additive). Media containing nucleobases/sides are labeled in bold font. The genes that are shown represent biosynthetic steps that could be bypassed or complemented by the exogenous nucleobases/sides in All64. Blank rows indicate that the gene was either essential or did not have detectable mutants at Time0 after recovery in rich media. Genes that were essential in all strains were excluded. Note that *purN* may be functionally redundant with *purT* except in *Bosea* OAE506, which lacks the latter; *purT* is not shown because it did not have a significant fitness score in any strain. Asterisks mark genes that were either essential, not detectable at Time0, or had a fitness score < –1 in All64.

We wondered if the importance of certain biosynthesis genes in the additive growers reflected a lack of transporters for the respective compounds. However, cross-referencing the RB-TnSeq results with our exometabolomics data suggested that this is not the case, as both additive growers fully consumed nearly all amino acids, as well as multiple nucleobases/sides, from All64 (Fig. 2). Such discrepancies between the ability to take up amino acids and nucleobases/sides and the ability to forgo the respective biosynthetic enzymes may indicate that the additive growers prioritized catabolizing those substrates instead of recycling them directly into biomass, though other factors such as timing, salvage capacity, and additional gene functions could also be relevant. Collectively, the RB-TnSeq results suggest that superadditive growth on All64 may arise in part from energy savings made possible by utilizing exogenous amino acids and nucleobases/sides as cellular building blocks, while additive growers appear to forego such savings and potentially experience other regulatory inefficiencies (Supplementary Notes).

### Biosynthetic protein abundance responds differently to mixed substrates across representative superadditive and additive growers

To gain additional insights into cellular resource allocation within superadditive versus additive strains, we performed proteomics to assess protein expression at the entry into stationary phase. We obtained high-quality data for five out of the six representative strains in all media that supported at least two doublings (superadditive *Lysobacter* OAE881 failed to yield usable data despite multiple attempts and robust growth). After confirming that the samples clustered by growth medium within each strain (Fig. 4a), we used differential protein expression analysis between medium conditions to determine 1) whether each strain displayed the capacity to consume multiple classes of substrates simultaneously in All64, and 2) whether the expression patterns for amino acid and nucleotide biosynthesis pathways were consistent with their fitness effects as observed by RB-TnSeq.

**Figure 4.**
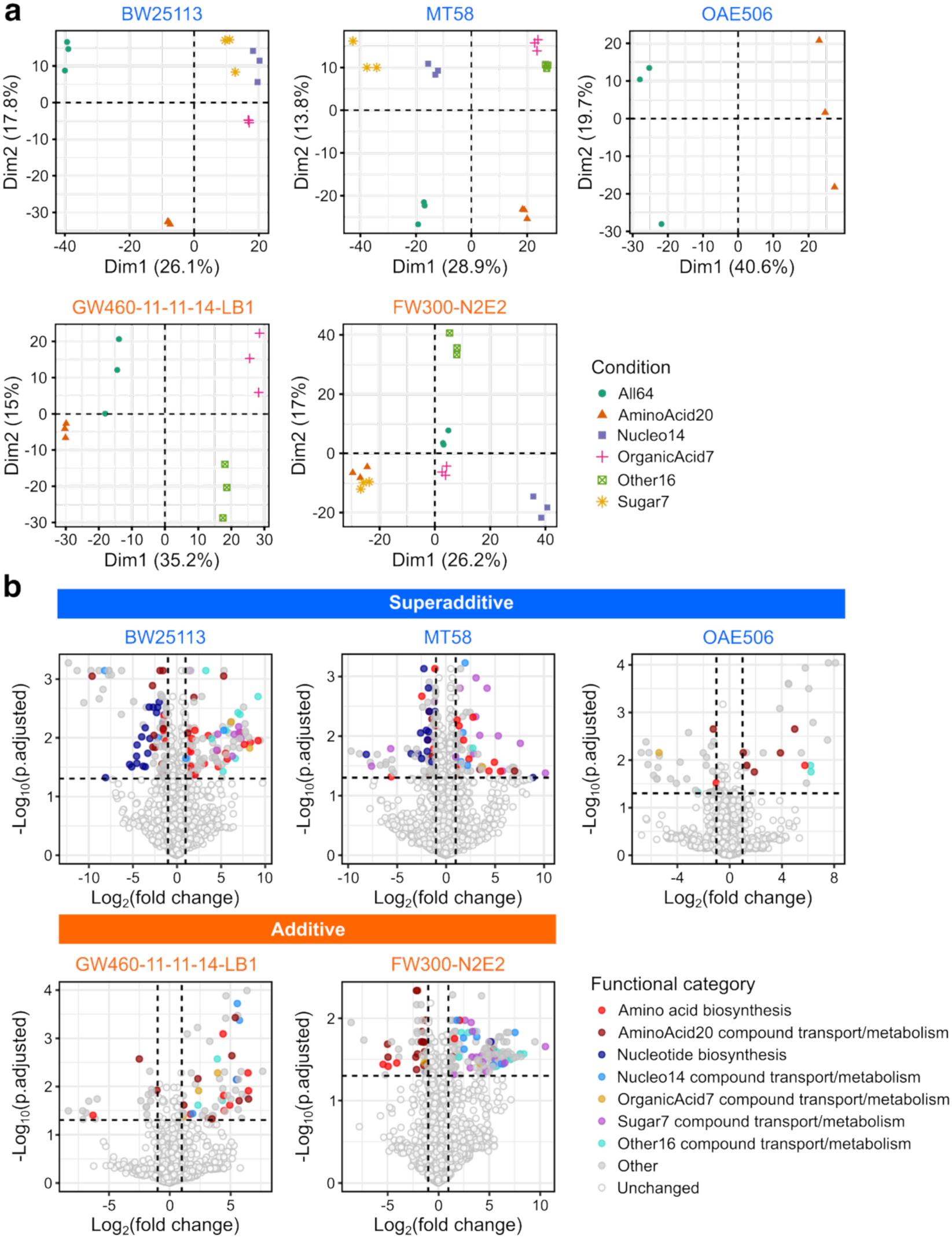
Protein expression patterns differ across representative superadditive and additive strains. **a)** PCA plots of the proteomics samples for each strain (n = 3 for each combination of strain and medium). **b)** Volcano plots highlighting differentially expressed proteins in All64 versus AminoAcid20 for each strain. Points are colored by the functional category to which the proteins belong. Dashed lines mark the cutoffs of log_2_ fold change of +/-1 and adjusted *p*-value < 0.05 (Welch’s t-test followed by the Benjamini-Hochberg procedure). Comparisons of All64 vs. other single-class media are shown in Extended Data Fig. 6.

For all strains, as expected, proteins related to consumption of compounds from multiple classes were upregulated in All64 relative to media lacking those compounds (Fig. 4b, Extended Data Fig. 6, Supplementary Table 7). All such compounds were indeed consumed by those strains in the exometabolomics experiment. Interestingly, however, not all compounds that were consumed in the exometabolomics experiment had upregulated transport or metabolic genes at the timepoint sampled for proteomics. Thus, the contemporaneous upregulation we observed for proteins related to multiple classes of carbon sources in All64 is potentially consistent with simultaneous consumption of those substrates, though verifying this would require time-resolved exometabolomics.

With regard to biosynthetic pathways, the expression patterns we observed were largely consistent with the RB-TnSeq results. Two out of the three superadditive growers downregulated numerous (8-20) nucleotide biosynthesis enzymes in All64 versus media without nucleobases/sides, whereas the additive growers downregulated at most one such enzyme (Fig. 4b, Extended Data Fig. 6, Supplementary Table 7). The same two superadditive growers also downregulated 10-28 amino acid biosynthesis enzymes in All64 compared to media without amino acids (Extended Data Fig. 6, Supplementary Table 7), whereas the additive growers downregulated relatively few (2-3 in most comparisons, though *P. fluorescens* FW300-N2E2 downregulated 9 in All64 vs. Other16 and 10 in All64 vs. Sugar7) (Extended Data Fig. 6, Supplementary Table 7). The low number of differentially expressed biosynthetic proteins for additive strains was not due to failure to detect such proteins, as the proteomics coverage of biosynthetic pathways was largely similar across all strains (Supplementary Table 8). The one exception to these trends was the superadditive grower *Bosea* OAE506, which did not downregulate any nucleotide biosynthetic enzymes in All64 versus AminoAcid20 (Fig. 4b). However, this strain’s regulation of amino acid biosynthesis enzymes could not be assessed as it only produced enough biomass for proteomics in media containing amino acids.

Overall, the downregulation of numerous biosynthetic enzymes by at least two of the three superadditive growers in All64 reinforces the idea that in some strains, superadditive growth may arise in part from energy savings conferred by salvaging cellular building blocks from the environment (including saved costs of biosynthetic enzyme expression), while the additive growers may have been hindered by constitutive expression of metabolically-expensive biosynthesis genes.

### Prediction of superadditive growth from genomic information

To facilitate future extension of the superadditive versus additive growth framework beyond our test strains, we sought to identify strategies for predicting these phenotypes from genotype. We first turned to a random forest classifier trained on a presence/absence matrix of KEGG orthologs^21^. This approach significantly outperformed two null models, with a median accuracy of 72.7% over 100 training/test splits (Fig. 5a). However, a phylogenomic nearest neighbor model performed equally well (Fig. 5a), raising the possibility that the random forest classifier was making predictions based on features correlated with phylogeny but not otherwise related to the phenotypes, as has been seen previously in other studies^22^. Indeed, most of the top ten most-important features identified by the random forest model do not have an obvious connection to a regulatory or biosynthetic function or metabolism of any carbon source present in All64 (Supplementary Table 9), nor did they have significant fitness effects in the RB-TnSeq experiments (if present in the tested strains). The one exception was K02564 (*nagB*, involved in catabolism of N-acetyl-glucosamine and N-acetyl-muramic acid), which was not only enriched in superadditive growers (Fig. 5b) but also required for full fitness of *E. coli* BW25113 and *Pantoea* MT58 in All64 (Supplementary Table 5), albeit this fitness effect was similar or stronger in Sugar7 and Other16 (which contain a higher concentration of N-acetyl-glucosamine or N-acetyl-muramic acid, respectively).

**Figure 5.**
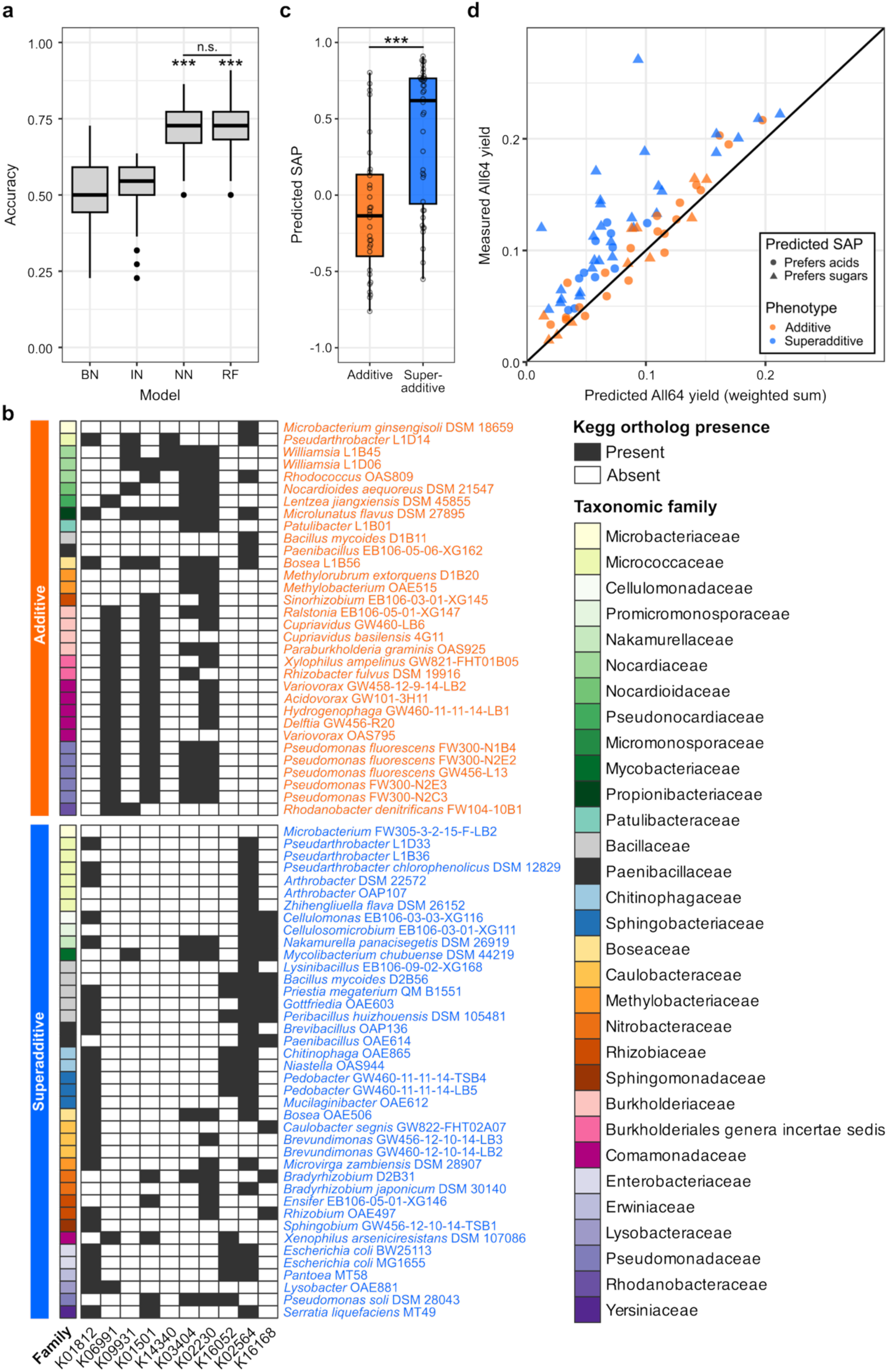
Three models achieve similar accuracy for predicting growth phenotype from genome content. **a)** Accuracy of different models for predicting superadditive vs. additive growth on All64 when the dataset was divided randomly 100 times into training and test sets (70%/30% split). BN = Bernoulli null, IN = identity null, NN = nearest neighbor (based on phylogenomic distance), RF = random forest (trained on KEGG ortholog presence/absence). \*\*\**p* < 0.001 (Mann-Whitney U test vs. each null model, followed by Benjamini-Hochberg procedure; the same significance level applies to the comparisons against both null models). **b)** Presence/absence matrix of the top ten most-important KEGG orthologs identified by the random forest classifier. Strains are organized by phenotype and taxonomic family. **c)** Distribution of predicted SAP (sugar-acid preference) values for strains belonging to the additive versus superadditive growth phenotypes. SAP was predicted from genome content following the procedure of Gralka et al., 2023. Negative values correspond to a preference for acids while positive values correspond to a preference for sugars. \*\*\**p* < 0.001 (Mann-Whitney U test). **d)** Plot of measured growth yield on All64 versus the weighted-sum prediction for the 72 genome-sequenced strains, colored by the assigned phenotype as in Fig. 1c (blue for superadditive, orange for additive) but with the point shape indicating whether each strain is predicted to prefer sugars (SAP > 0, triangles) or acids (SAP < 0, circles).

We next considered whether superadditive versus additive growth could be related to genomically-encoded preferences for different types of carbon sources^23^. In particular, we hypothesized that superadditive strains might tend to prioritize catabolizing glycolytic substrates (sugars), which could allow simultaneously imported amino acids to be funneled directly into translation, facilitating the downregulation of costly biosynthetic pathways. Conversely, strains that prefer gluconeogenic substrates (amino acids and organic acids) might prioritize degrading imported amino acids for carbon and energy over using them to make proteins, at the cost of maintaining a higher biosynthetic capacity to keep up with translational needs. Indeed, predicted sugar-acid preference (SAP)^23^ was significantly higher on average among superadditive growers (median SAP = 0.62) than additive growers (median SAP = −0.14) (Fig. 5c). Empirically, superadditive growers also tended to have higher growth rates and yields on Sugar7 compared to additive growers, and vice versa on AminoAcid20 and OrganicAcid7, although these differences were not statistically significant (Extended Data Fig. 7). Simply predicting superadditive growth for sugar-preferring strains (predicted SAP > 0) and additive growth for acid-preferring strains (predicted SAP < 0) resulted in 69% accuracy (Fig. 5d), though as with the random forest model, this result may partly reflect lineage structure.

Finally, to explore whether a more granular mechanistic approach to utilizing genomic information would provide additional explanatory power, or better predict superadditive growth, we took advantage of genome-scale metabolic models (GEMs) that were recently reconstructed for a subset of our tested strains^24^. Using Computation Of Microbial Ecosystems in Time and Space (COMETS)^25^ to perform dynamic flux balance analysis (dFBA), we simulated growth to stationary phase in our six experimental media. Other than setting environmental metabolite abundance in COMETS to mirror the experimental media composition, models were used as previously published^24^, assuming COMETS default uptake kinetic parameters^25^ uniformly across metabolites and strains. Even without uptake parameter fitting, model-predicted growth yields across strains recapitulated some of the trends observed in the empirical data (Extended Data Fig. 8a). Moreover, as shown in Extended Data Fig. 8b, COMETS predicts modest superadditivity for several strains, in line with previous FBA calculations suggesting that many carbon sources exhibit cooperativity in promoting growth^14^. Interestingly, COMETS also predicted that a higher fraction of amino acid uptake would be used directly as building blocks for biomass in All64 versus AminoAcid20 (Extended Data Fig. 8c). However, COMETS consistently underestimated the relative biomass gain on All64 versus the weighted sum for superadditive strains (Extended Data Fig. 8d) and could not distinguish between empirically superadditive versus additive strains (Extended Data Fig. 8e). These limitations may be attributed to the lack of metabolite-specific uptake kinetic parameters, or to the fact that standard genome models lack a detailed regulatory layer that could account for individual strains capitalizing on or foregoing the savings induced by multiple building block availability.

For the same strains, the random forest model correctly predicted the growth phenotype >90% of the time for 10 out of 14 strains, and >55% of the time for all but two strains (*Bosea* OAE506, a superadditive grower, and *Methylobacterium* OAE515, an additive grower). These results suggest that while GEMs are capable of reproducing superadditivity^26^, without further uptake parameter fitting they fail to capture the full range of deviations from additive growth or to accurately distinguish between additive and superadditive strains. By contrast, a relatively simple random forest classifier achieved significantly better-than-random binary prediction of superadditive versus additive growth, while phylogeny and SAP represent similarly useful, if still imperfect, heuristics, though generalizability beyond our strain collection remains to be determined. This contrast highlights the tradeoff between data-driven higher-accuracy machine learning-based approaches and mechanistic approaches, whose accuracy is limited by the amount of knowledge encoded, but whose detailed representation of biological entities provides explanatory power, potentially pointing to the need of future hybrid approaches.

## Discussion

In this study, we have found that superadditive growth on a dilute, carbon-limited mixture of carbon sources representing different functional classes is a common, though not universal, phenomenon in heterotrophic bacteria. By using defined media and pursuing a multimodal mechanistic investigation, we have identified multiple factors that likely contribute to superadditive growth. Specifically, superadditive growers consumed numerous substrates to a greater degree in the mixed-class medium than in single-class media, and were less dependent on and often downregulated amino acid and nucleobase/side biosynthesis genes when those compounds were provided exogenously (Supplementary Discussion). Conversely, additive growers consumed the vast majority of substrates to a similar extent in both the mixed-class and single-class media, were obligately dependent on biosynthesis of several amino acids and nucleobases/sides, and failed to downregulate most biosynthetic genes in the mixed-class medium compared to media lacking amino acids or nucleobases/sides. We have also shown that superadditive versus additive growth under our experimental conditions can be predicted from genome content with ∼70% accuracy, laying the groundwork for assessing the putative prevalence of these phenotypes across environmental metagenomic datasets.

Notably, superadditive versus additive growth is an axis of variation in bacteria that is potentially complementary to other frameworks for physiological classification of heterotrophic bacteria, such as copiotrophs versus oligotrophs^27^ and metabolic generalists versus specialists^28,29^. Many if not most of the strains we tested are copiotrophs, yet they spanned a spectrum from clearly additive to exceptionally superadditive. Moreover, superadditive and additive strains alike depleted a wide range of compounds from All64, suggesting that these phenotypes may be largely unrelated to the breadth of a strain’s metabolic niche. We therefore propose that adding the dimension of superadditive versus additive growth to these existing frameworks may help improve the predictive understanding of how different strains, and ultimately communities, respond to changes in nutrient diversity, which remains a significant challenge^2,29,30^.

Given that our definition of superadditive growth is based on within-strain comparisons (across media conditions), we cannot yet make generalized predictions for how the balance of superadditive versus additive growers within microbial communities may affect phenomena such as inter-strain competition or soil carbon turnover dynamics (Supplementary Discussion). However, another recent study similarly reported that diverse bacterial taxa exhibit enhanced carbon consumption when grown on a mixture of 22 metabolites versus single carbon sources^30^. Intriguingly, the authors observed that communities containing such strains experienced stronger competitive interactions with increasing nutrient diversity, causing biodiversity losses. However, they did not explicitly compare the growth yields of individual strains on mixed carbon sources to the additive null expectation. Complementing this earlier work, we have observed greatly enhanced carbon consumption when different classes of substrates are combined is characteristic of superadditive growers. We therefore speculate that communities enriched in superadditive growers could likewise experience intensified competition with increasing nutrient diversity, although testing this hypothesis will require further experimental work. Moreover, the physiological differences between superadditive and additive growers may also manifest in ways beyond growth yield on mixtures of diverse carbon sources, with added potential to impact interspecies interactions and community structure (Supplementary Discussion).

Finally, while we have identified differences between superadditive versus additive growers with respect to carbon uptake and conditional essentiality of biosynthetic pathways, these findings are based on a relatively small subset of strains from Pseudomonadota, which were chosen based on the availability of existing RB-TnSeq libraries. Follow-up studies will be needed to assess the conservation of these patterns across a broader range of taxa representing each phenotype. Moreover, factors such as concentration-dependent utilization, differences in growth timing, and other physiological constraints may also contribute to these growth phenotypes, and the underlying biochemical mechanisms at the molecular level remain to be elucidated (Supplementary Discussion).

Taken together, by formalizing the concept of superadditive versus additive growth, uncovering the prevalence of these phenotypes across diverse bacteria, and highlighting some of the key underlying physiological differences, our work adds a new dimension to the understanding of bacterial growth on mixed substrates. Importantly, the prevalence of superadditive growth and superadditive carbon source consumption across diverse taxa, with the latter being confirmed by an independent study, suggests that while single-carbon-source growth assays have been a mainstay of attempts to characterize the metabolic niches of heterotrophic bacteria^23,29,30^, this approach may frequently underestimate the realized niche under naturalistic conditions. Such findings underscore that for studies seeking to understand microbial ecology in terms of niche theory, it is critical to use multiomics assays to empirically define realized metabolic niches in the environment of interest^28^. Future work toward resolving the molecular basis of superadditive versus additive growth across diverse taxa will not only help to improve the accuracy of genotype-to-phenotype predictions, but could also inform synthetic biology efforts and the refinement of genome-scale metabolic models, which are useful for a variety of applications. Additionally, assessing the environmental distribution of these phenotypes, whether through culture-based or metagenomic surveys, could help unveil the selective pressures and evolutionary forces shaping this underrecognized yet potentially broadly relevant axis of diversification among heterotrophic bacteria.

## Methods

### Bacterial strains tested in this study

We selected 100 diverse bacterial strains (Supplementary Table 1), including 25 type strains (initially isolated from soil; sourced from DSMZ, Germany) representing globally dominant soil bacteria^31^, 73 isolates from various soil environments (biocrusts, switchgrass rhizosphere, agricultural fields, sediments, and other soils), and two model *E. coli* strains— MG1655, the wild type K-12 strain, and BW25113, a K-12 strain and parent of the Keio Collection^32^ that has an established genome-wide, randomly barcoded transposon mutant library^17^. In total, this list of 100 bacterial strains spans four phyla, nine classes, 21 orders, and 37 families.

### Media conditions

To compare how different bacteria vary in their growth in simple vs. complex mixtures of substrates, we set up six different defined media (Supplementary Table 2) to grow each individual strain. To construct the defined media, we chose a total of 64 representative metabolites (carbon sources) belonging to a variety of chemical classes (e.g., sugar, amino acid, organic acid, etc.), that were previously described to be representative of soil organic carbon profiles and support a wide range of bacterial growth as a defined medium^15^. Across the six media, the total molar concentration of metabolites was held constant (2560 µM), but the number and type of metabolites varied. Specifically, we had five simpler but more concentrated media, four of which each contain a mixture of metabolites from a single chemical class (Sugar7, OrganicAcid7, AminoAcid20, Nucleo14), while the fifth, Other16, was a catch-all for compounds that do not clearly belong to the other classes. For example, AminoAcid20 means a mixture of 20 different amino acid compounds each at an equal concentration of 128 µM. In addition, we also had a more complex but diluted medium containing all 64 metabolites (All64) each at an equal concentration of 40 µM. In all media, metabolite compounds were mixed into a base medium containing KH_2_PO_4_ (0.6 g L^−1^) and NH_4_Cl (1.5 g L^−1^) to provide excess inorganic nitrogen and phosphorus, along with Wolfe’s vitamins and Wolfe’s mineral solutions each at 1x concentration, as in defined media used previously for culturing a variety of bacterial isolates^33^. The final pH was adjusted to 7.2 and the media were filtered through a 0.22 μm PES membrane to sterilize. All media components were purchased from Sigma-Aldrich, except for D-(+)-trehalose dihydrate (Santa Cruz Biotechnology), xylose (Spectrum Chemical), and sn-glycero-3-phosphocholine (Cayman Chemical).

We chose to standardize the total molar (molecule) concentration rather than total carbon content for several reasons. First, our primary aim was to compare growth on chemically simple but concentrated pools of substrates vs. a chemically diverse but dilute pool. Holding molar concentration constant ensures that the summed molar contribution of the five single class media equals that of the complex All64 medium, making the weighted-sum prediction of yield directly interpretable (i.e., each single class medium represents a defined molar fraction of the complete mixture). Second, standardizing by carbon would have required unequal molar concentrations across classes because carbon content per molecule varies widely (Supplementary Table 2; e.g., amino acid alanine having only 3 carbon atoms vs. nucleoside inosine having 10 carbon atoms). This would significantly confound substrate concentration, which is a key determinant of uptake kinetics and osmotic conditions, with substrate identity. By fixing molar concentration, each metabolite is present at a defined, comparable per-molecule concentration, and any differences in growth can be attributed to substrate identity and diversity rather than to differing numbers of dissolved molecules.

We acknowledge that this design results in differing total carbon and nitrogen contents across media, which we calculated exactly for each medium (Supplementary Table 2). These differences are relevant primarily when directly comparing absolute growth yields across medium types. Importantly, however, they do not affect our central analysis: because the weighted-sum model (Equation 1, see below) compares the measured All64 yield against a prediction built from the same metabolites at the same per-molecule concentrations, any C (or N) contributed by a given metabolite is accounted for identically on both sides of the comparison. Our test for superadditive vs. additive growth is therefore internally consistent with respect to carbon content.

### Growth profiling in single-class vs. mixed-class media

To characterize the growth of individual strains in each of the six different media, we revived each isolate from freezer glycerol stocks onto Reasoner’s 2A (R2A)^34^ agar plates and incubated at 27 °C. Single colonies were picked and inoculated into R2A broth and incubated at 27 °C with shaking at 200 rpm as the starter liquid culture. Each starter culture was then centrifuged, cell pellets washed with fresh base medium (pH 7.2), and resuspended in each respective medium to a final optical density at 600 nm (OD_600_) of 0.05 to initiate the growth experiment. Growth was tracked by taking OD_600_ readings every 30 min with shaking at 27 °C for 64 h using a Biotek Synergy HT microplate reader (Agilent Technologies, Santa Clara, CA). Respective uninoculated media were included as an abiotic control. At least two, and in most cases three or more independent replicate growth curves were done on different days using freshly made media to improve the reproducibility of the results. Growth curves were generated by normalizing strain OD_600_ readings against the respective abiotic control readings over time. The data were smoothed to mitigate noisiness using the moving-median function from the *gcplyr* package^35^ in R, with a window of 9 time points, and the maximum OD_600_ after smoothing was calculated as a proxy for yield. We chose the median as the summary statistic for the yield across replicates due to occasional outlier results that would have skewed the means for some strains, and correspondingly used median absolute deviation (MAD) from the median as the metric of variability. Growth rates (shown in Extended Data Fig. 7b) were inferred by using the *growthcurver* package^36^ in R to fit a logistic growth model.

Of the 100 strains, eight failed to grow in starter culture and 16 failed to grow in any of the six defined media, leaving 76 strains for downstream analysis. For these remaining 76 strains, we used a weighted-sum model to predict their yield grown on the diluted mixture of 64 metabolites (All64) from yield measured on the five individual fractional metabolite classes, and compared this with the actual measured yield in the All64 mixture.

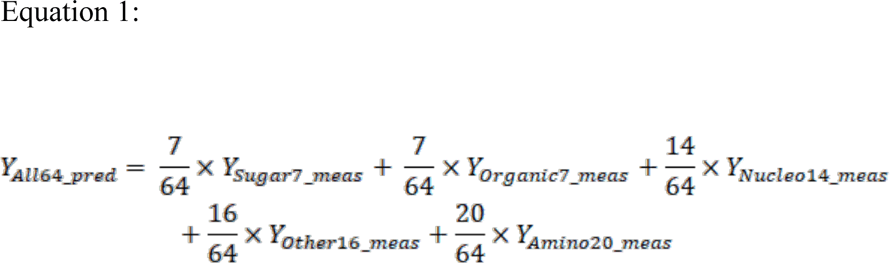

Where “Y” represents yield, “pred” represents the weighted-sum predicted yield, and “meas” represents the measured yield. This is a weighted-sum model because each single class medium contributes to the prediction in proportion to the number of metabolites in that class, i.e., the fraction of the 64-metabolite mixture that it represents (e.g., 7/64 for Sugar7, 20/64 for AminoAcid20). Because the total molar concentration was held constant across all media, weighting reconstructs the All64 substrate amounts under the assumption that growth yield scales linearly with concentration, so weighting by class size reconstructs the expected yield if each metabolite class contributed independently and additively to growth. A measured All64 yield that exceeds this prediction by our screening criterion, as described below, therefore indicates superadditive growth (the diverse mixture supports more growth than the sum of its parts), whereas failure to meet this criterion indicates that superadditivity was not detected.

### Classification of strains as superadditive versus additive growers

When seeking to compare yields on All64 versus the weighted sum prediction while accounting for the variability among replicates, one difficulty we faced is that there is no set formula for propagating error based on MAD when adding two or more variables. Therefore, to approximate a metric of uncertainty for the weighted sum estimate based on the variability in the underlying data, here MAD was treated as a physical measurement uncertainty, i.e. the uncertainty estimate of the weighted sum was taken as the weighted sum of the MAD for each variable. This is a very conservative approach: when we repeatedly simulated data for five normal distributions (a, b, c, d, e), where the mean and standard deviation were randomly sampled within the range of median and scaled MAD values we observed for the five single-class media, MAD(a) + MAD(b) + MAD(c) + MAD(d) + MAD(e) was always greater than MAD(a + b + c + d + e).

We then defined superadditive growth as All64 yield - All64 MAD (scaled by 1.4826) greater than the weighted sum prediction + weighted sum estimated error (again scaled by 1.4826) based on the following reasoning. Assuming that the underlying distribution of growth yields for a given strain x medium combination is normal (likely a reasonable assumption as this is true for many continuous biological variables that are influenced by numerous additive independent factors), 68% of data points should be within 1 standard deviation (sd, equivalent to 1.4826 MAD) of the median. Under this assumption, our cutoff means that <16% of measurements of All64 growth yield (if we could take a very large number of measurements) would be expected to overlap with the range of values of the corresponding weighted sum prediction (again based on a very large number of measurements of growth on the other five media) and vice versa. When t-tests are simulated on randomly-generated summary statistics for normal distributions within the range of values we observed for growth yield, where mean2 = mean1 + sd1 + sd2 (essentially simulating data right at our cutoff), the Benjamini-Hochberg-adjusted p-values are < 0.05 the vast majority of the time for 6+ replicates per condition, and a majority of the time for 5 replicates. For this to be true with fewer replicates, e.g. 3, the difference between mean1 and mean2 has to be much greater (e.g. 2*sd1 + 2*sd2), but considering that our approach to estimating the uncertainty in the weighted sum prediction almost certainly overestimates the true uncertainty, we decided to proceed with the less aggressive cutoff. Even so, note that some of the strains classified as additive due to high variability may not be truly additive but rather “not detectably superadditive” with the current number of replicates. Finally, one remaining caveat is that since we used OD_600_ as our metric for growth yield, our analysis is predicated on a consistent relationship between OD_600_ and biomass for each strain across the different media.

### Phylogenomic tree construction

Genomes were obtained for 72 out of the 76 phenotyped strains from the sources described in Supplementary Table 1. The remaining four strains have not been fully sequenced. We constructed an unrooted phylogenomic tree for the 72 strains with GToTree^37^ using default settings for bacteria, with the amino acid FASTA files for all genomes as the input. The resulting Newick tree file was visualized using the *ggtreeExtra* package in R.

### Exometabolomic profiling of spent growth media

To explore potential mechanisms of superadditive growth, we selected six representative strains from the superadditive and additive groups, prioritizing those for which mutant libraries were available for further genetic examination (as described below). First, we examined how substrate utilization differs in the diluted mixture of 64 diverse substrates (All64) vs. higher concentrations of the five individual classes of substrates that comprise the mixture (i.e., the single-class media). Starter cultures of the six strains were prepared in the same manner as for growth profiling. Afterwards, cell pellets were collected, washed with fresh base medium (pH 7.2), and resuspended in 25 mL of each respective growth medium to an initial OD_600_ of 0.02 to initiate the growth experiment in sterile PETG flasks (Nest Scientific #781011) at 27 °C with 200 rpm shaking. We first monitored growth of each strain in flasks over time and compared the resulting curves with the plate reader results to establish the expected growth trajectory for each strain and medium combination. During the sampling experiment, we therefore had an a priori expectation of the growth pattern but still periodically sampled each culture to confirm its growth stage. There were three replicate flasks for each strain and medium combination. Three replicates for each uninoculated medium were included as an abiotic treatment control. Each strain and medium combination was sampled at late exponential/early stationary phase, together with the respective uninoculated medium control, by centrifugation at 5000 x g for 10 min at 4 °C. Supernatants (1.5 mL) were collected into 2 mL tubes and stored at −80 °C until exometabolomics analysis. Cell pellets were frozen at −80 °C for proteomics analysis as described below.

For exometabolomics analysis, the frozen supernatant samples were lyophilized and resuspended in 150 μL of LCMS-grade methanol spiked with a labeled internal standard mix (Supplementary Table 4). The resuspended samples were vortexed, sonicated in an ice water bath, and centrifuged. The resulting supernatants were filtered through 0.2 μm modified nylon membrane centrifugal filters, and filtrates were collected for analysis. Metabolites were separated via hydrophilic interaction liquid chromatography (HILIC) and detected with high resolution mass spectrometry. Specifically, the samples were analyzed using an InfinityLab Poroshell 120 HILIC-Z column (Agilent) on an Agilent 1290 UHPLC stack connected to an Orbitrap Exploris 120 mass spectrometer equipped with a Heated Electrospray Ionization (HESI) source probe. Separation via gradient elution was followed by MS1 and data-dependent MS2 collection (top two most-abundant MS1 ions not previously fragmented in the last seven seconds). Detailed instrument parameters and LC conditions are described in Supplementary Table 4.

Metabolites were identified through a targeted analysis pipeline using Metabolite Atlas^38^ (https://github.com/biorack/metatlas), whereby m/z, retention time, and fragmentation spectra were compared to a library of analytical standards previously analyzed under the same conditions. In total, 63 out of the 64 metabolites were identified, except spermidine, which was not detected in any sample including the uninoculated controls. The raw metabolite abundances were calculated as peak heights for most compounds, or peak areas for compounds that eluted as multiple peaks (Supplementary Table 4). For compounds with labeled internal standards (Supplementary Table 4), the peak heights or peak areas of the unlabeled compounds were normalized relative to the internal standard peak heights or areas. The ratio of metabolite abundances between the spent supernatant and uninoculated medium control was calculated and multiplied by 100 to obtain the percent remaining of each carbon source in the supernatants relative to the starting concentration. A ratio >100% indicates net excretion while a ratio <100% indicates net depletion of that metabolite from the medium. To calculate absolute net consumption/excretion for each carbon source, the fraction remaining was multiplied by its known starting concentration in the respective medium. For each strain, to identify significant differences in either the percent remaining or absolute net consumption of each carbon source between All64 and the respective single-class medium (n = 3), we employed Welch’s t-test followed by the Benjamini-Hochberg procedure to control the false discovery rate. We considered an adjusted *p*-value < 0.05 to be significant. On top of this, if a compound met any of the following criteria, we did not consider it to exhibit meaningfully increased consumption in All64 even if the adjusted *p*-values were <0.05: 1) compounds with net excretion in both All64 and the single-class medium; 2) compounds with a difference of <5% between the percent remaining in All64 versus the single-class medium; 3) compounds for which the percent remaining in All64 was not significantly different from 100% (two-sided one-sample t-test).

### Competitive mutant fitness assays using RB-TnSeq

To investigate the genetic factors underlying the differential growth and substrate metabolism of superadditive and additive strains in the diluted mixture of All64 vs. concentrated single class of substrates, we leveraged the available RB-TnSeq^17^ libraries that have been developed for the six strains we focused on. The libraries for *E. coli* BW25113, *Pantoea* sp. MT58, *Lysobacter* sp. OAE881, and *Pseudomonas fluorescens* FW300-N2E2 have been previously described^17–20^, while the libraries for *Bosea* sp. OAE506 and *Hydrogenophaga* sp. GW460-11-11-14-LB1 were generated prior to this study but are being reported here for the first time.

The *Bosea* sp. OAE506 library was made by conjugating the wild-type parent strain with *E. coli* donor strain WM3064 carrying the barcoded mariner transposon delivery vector pHLL250_NN1 (strain AMD290). *Bosea* sp. OAE506 was pregrown at 30 °C in R2A, and AMD290 was pregrown in lysogeny broth (LB) + carbenicillin (50 µg/mL) + diaminopimelic acid (DAP, 300 µM) at 37 °C. Conjugation was performed at 30 °C on LB + DAP plates (without a filter) for 5 hours. Mutants were selected on R2A + kanamycin (50 µg/mL) plates at 30 °C and pooled as previously described to make the final RB-TnSeq library (Bosea_OAE506_ML2). The *Hydrogenophaga* sp. GW460-11-11-14-LB1 library was likewise made via conjugation from *E. coli* donor strain AMD290. *Hydrogenophaga* sp. GW460-11-11-14-LB1 was pregrown at 30 °C in LB, and AMD290 was pregrown in LB + Carb50 + DAP at 37 °C. Conjugation was performed at 30 °C on LB + DAP plates (without a filter) for 12 hours. Mutants were selected on LB + kanamycin (25 µg/mL) plates at 30 °C and pooled as previously described to make the final RB-TnSeq library (Phaga5_ML11). For both libraries, the transposon insertions were mapped and associated with the random DNA barcodes according to a previously described protocol^17^. The Bosea_OAE506_ML2 library comprises 135,883 mapped mutants, while Phaga5_ML11 comprises 138,294 mapped mutants.

For the pooled mutant fitness assays, a single aliquot of each mutant library was thawed and revived in fresh rich medium (LB for *E. coli* BW25113, *Pantoea* sp. MT58, *Pseudomonas fluorescens* FW300-N2E2; R2A for *Lysobacter* sp. OAE881, *Bosea* sp. OAE506, and *Hydrogenophaga* sp. GW460-11-11-14-LB1) supplemented with 25 µg mL^−1^ kanamycin. These starter cultures were grown overnight to mid-exponential phase, then sampled to measure OD_600_, and cell pellets were collected as a Time0 reference. Then, an appropriate volume of culture was centrifuged, and the pellets were washed twice with fresh base medium (pH 7.2) and resuspended in 15 mL of each respective medium to a starting OD_600_ of 0.01 in sterile culture tubes. The cultures were incubated at 27 °C with shaking at 200 rpm. There were three replicate tubes for each strain and medium combination. When the mutant library cultures reached saturated growth, we measured OD_600_ and collected two pellet aliquots: one stored at −80 °C and one used to inoculate fresh medium at the same starting OD_600_ to begin the next growth cycle. This process was repeated twice such that the final samples were collected after the third successive growth cycle. This allowed mutants with subtle growth advantages or disadvantages in a given medium to be enriched or depleted to a greater extent, improving our ability to detect genes that relate to the differential growth and substrate metabolism across the different media. All cell pellets were stored at −80 °C prior to genomic DNA extraction when all mutant assays were completed.

Genomic DNA was extracted in 96-well format using a QIAamp 96 DNA QIAcube HT kit (Qiagen #51331) following the manufacturer’s instructions, except that a vacuum manifold was used for processing the samples. BarSeq was performed as previously described^18^ using unique dual indexed primers (https://bitbucket.org/berkeleylab/feba/src/master/primers/). BarSeq samples were sequenced on the NovaSeq X 10B platform (1×100 bp). The fitness of each unique barcoded mutant was calculated as the binary logarithm of the ratio of its final relative abundance to its Time0 relative abundance. Gene fitness values and a *t*-like statistic for assessing significance were subsequently calculated as previously described^17^.

For the analysis of amino acid biosynthesis pathways, for each strain we determined the most likely pathways and constituent genes using GapMind for amino acid biosynthesis^39^. The specific genes assigned to each pathway are documented in Supplementary Table 6. We determined gene essentiality from TnSeq data according to previously published criteria^18^. Genes that did not qualify as essential based on the TnSeq results were still considered highly important if no barcodes for insertions in that gene were detected at Time0 of the BarSeq experiments. We considered a pathway to be essential if all predicted members of the pathway were essential/missing at Time0 except for genes that showed no differential fitness effects between conditions with and without amino acids, as the latter is a sign of likely functional redundancy. Alanine and glycine were excluded from Fig. 3a because in all strains, the relevant genes were either functionally redundant, or essential but known to also be required for other biosynthetic processes. For each non-essential pathway, we calculated the mean fitness score across all pathway members after excluding any essential genes that were part of the pathway (most such genes were those known to also be required for other biosynthetic processes, such as biosynthesis of cofactors or peptidoglycan). We chose to aggregate the fitness values at the pathway level since not all strains shared the same biosynthetic pathways for each amino acid, complicating gene-level comparisons.

For the analysis of nucleotide biosynthesis genes, we focused on genes encoding enzymatic steps that could be complemented by the nucleobases/sides provided in All64 and Nucleo14. Reassuringly, these were the genes for which mutants in *E. coli* BW25113 were fully rescued by All64 and Nucleo14. Unlike amino acid biosynthesis, all strains shared the same biosynthetic steps for nucleotides, enabling direct gene-level comparisons of fitness effects.

For the analysis of genes with positive fitness scores specific to All64, we focused on genes with |t| > 4 and fitness score > 1 in All64, with fitness scores < 0.5 in all other conditions for the same strain. For the analysis of genes related to catabolism of different carbon sources, we focused on genes with |t| > 4 and fitness scores < −1 and verified the likely functions of such genes by cross-referencing the genome annotations with BioCyc^40^.

### Proteomic analysis

To assess how the superadditive vs. additive strains differentially regulate their proteomes when grown in the diluted mixture of All64 vs. concentrated single classes of substrates, we performed comparative proteomics. Cell pellets were collected at early stationary phase either from the exometabolomic profiling experiment described above (*E. coli* BW25113 and *Pantoea* MT58) or in a separate experiment performed according to the same procedure (*Bosea* OAE506, *Hydrogenophaga* GW460-11-11-14-LB1, *Pseudomonas fluorescens* FW300-N2E2). Each pellet comprised 1-2 OD_600_ units of biomass. Protein was extracted from cell pellets and tryptic peptides were prepared by following established proteomic sample preparation protocol^41^. Briefly, cell pellets were resuspended in Qiagen P2 Lysis Buffer (Qiagen, Germany) to promote cell lysis. Proteins were precipitated with addition of 1 mM NaCl and 4 x vol acetone, followed by two additional washes with 80% acetone in water. The recovered protein pellet was homogenized by pipetting mixing with 100 mM ammonium bicarbonate in 20% methanol. Protein concentration was determined by the DC protein assay (BioRad, USA). Protein reduction was accomplished using 5 mM tris 2-(carboxyethyl)phosphine (TCEP) for 30 min at room temperature, and alkylation was performed with 10 mM iodoacetamide (IAM; final concentration) for 30 min at room temperature in the dark. Overnight digestion with trypsin was accomplished with a 1:50 trypsin:total protein ratio. The resulting peptide samples were analyzed on an Agilent 1290 UHPLC system coupled to a Thermo Scientific Orbitrap Exploris 480 mass spectrometer for discovery proteomics^42^. Briefly, peptide samples were loaded onto an Ascentis® ES-C18 Column (Sigma-Aldrich, USA) and were eluted from the column by using a 10 minute gradient from 98% solvent A (0.1 % FA in H2O) and 2% solvent B (0.1% FA in ACN) to 65% solvent A and 35% solvent B. Eluting peptides were introduced to the mass spectrometer operating in positive-ion mode and were measured in data-independent acquisition (DIA) mode with a duty cycle of 3 survey scans from m/z 380 to m/z 985 and 45 tandem mass spectrometry (MS2) scans with precursor isolation width of 13.5 m/z to cover the mass range.

DIA raw data files were analyzed by an integrated software suite, DIA-NN^43^. The databases used in the DIA-NN search (library-free mode) are the proteome FASTA sequences acquired from the Fitness Browser (https://fit.genomics.lbl.gov/) for the strains under investigation, plus common proteomic contaminants. DIA-NN determines mass tolerances automatically based on first pass analysis of the samples with automated determination of optimal mass accuracies. The retention time extraction window was determined individually for all MS runs analyzed via the automated optimization procedure implemented in DIA-NN. Protein inference was enabled, and the quantification strategy was set to Robust LC = High Accuracy. Output main DIA-NN reports were filtered with a global false discovery rate set at 0.01 (FDR <= 0.01) on both the precursor level and protein group level. The sum peak area of all peptides from an identified protein was used for quantitative analysis. To compare the proteomes, we calculated the fold change of protein abundances for each strain in All64 vs. each single-class media. To assess statistical significance, we used Welch’s *t*-test (n = 3) on log_2_-transformed abundances followed by the Benjamini-Hochberg procedure for controlling the false discovery rate, yielding adjusted *p*-values. Proteins with an adjusted *p*-value < 0.05 and log_2_ fold change > 1 (upregulated) or < −1 (downregulated) were considered to be significantly altered. Differentially expressed proteins were manually assigned to relevant functional classes (amino acid biosynthesis, nucleotide biosynthesis, or metabolism of the different classes of carbon sources in All64) by cross-referencing the gene names and/or gene product descriptions from the annotated genomes with BioCyc^40^.

### Genotype-to-phenotype predictions

Genomes were annotated with Kegg orthologs (KOs) using eggNOG Mapper^44^ with eggNOG Database 5.0.2 and default settings on the European Galaxy server^45^. In instances where eggNOG Mapper returned more than one KO match for a gene, we kept only the first match. We then generated a presence/absence matrix of all detected KOs across all 72 genome-sequenced strains. We excluded KOs that appeared in only one strain or in all strains. We split the dataset 100 times randomly into training sets (70%) and test sets (30%) and trained a random forest classifier on the KO presence/absence matrix using the *ranger* package^46^ in R, with mtry set to the maximum (equal to the total number of features). We used the permutation method in *ranger* to calculate feature importance.

For the phylogenomic nearest neighbor model, we again split the data set into 100 training/test sets (70%/30%) with the same partitions as above. Test strains were assigned the phenotype of their closest relative, based on phylogenomic distance according to the phylogenomic tree shown in Extended Data Fig. 2. For the identity null model, the data set was split into the same 100 training/test sets (70%/30%) and for each iteration, all test strains were assigned the most prevalent phenotype in the respective training set. For all of the aforementioned models, accuracy was calculated as the fraction of test strains for which the predicted phenotype matched their empirical phenotype. For the Bernoulli null model, the phenotype was guessed for each strain with the same probability as the occurrence of each phenotype across the entire dataset. In other words, this model essentially performs a weighted coin flip. This procedure was repeated 100 times for each strain.

For calculation of predicted sugar-acid preference, we followed the procedure described by Gralka et al., 2023^23^. Briefly, we calculated the relative abundance in each genome of KOs assigned to either sugar metabolism or acid metabolism as defined by Gralka et al. These relative abundances were used as the input to the formula described by Gralka et al. to transform the ratio of sugar-related KO relative abundance and acid-related KO relative abundance into a score ranging from −1 to 1, where values < 0 indicate a preference for acids and values > 0 indicate a preference for sugars.

To predict biomass yield on different media compositions using GEMs, we used COMETS version 2.12.4^25^, cometspy version 0.5.2, and the Gurobi solver version 10.0.1. Default biomass objectives were maintained for these GEMs. Simulations were run for 144 steps of 0.5 hours each (72 hours total), to ensure all strains grow to stationary phase. Space width and media concentrations as defined earlier in the study were set in to simulate 200 µL test tubes, with a space width of 0.585 cm in a single well-mixed 1×1 grid. Death and dilution rates were set to 0. Vmax and Km were set as modeling assumptions to 10 mmol / (gDW * h) and 10^−6^ M respectively for all metabolites to allow for complete uptake of media. Unless otherwise stated, parameters were set to be the default for the stated version of COMETS. Initial biomass was set to 10^−6^ g dry weight. To determine the fate of amino acids taken up from the medium, we reproduced the first COMETS time step for each model and medium in cobrapy (version 0.32.1, GLPK solver) and solved using pFBA. For each amino acid, the fraction of uptake incorporated into biomass was calculated as min(B, U)/U, where U is net uptake from the medium and B is the flux consumed by the biomass reaction.

## Supporting information

Supplementary Discussion

## Data availability

The raw exometabolomics data are available at the Global Natural Products Social Molecular Networking repository (https://gnps2.org/status?task=c8eba28fa9b7476ea343c13488d49b3e). RB-TnSeq fitness data are available at the Fitness Browser website (https://fit.genomics.lbl.gov). Genome repository locations and accession numbers are listed in Supplementary Table 1. The generated mass spectrometry proteomics data have been deposited to the ProteomeXchange Consortium via the PRIDE partner repository^48^ with the dataset identifier PXD084911.

## Code availability

The code for the Metabolite Atlas pipeline is available at https://github.com/biorack/metatlas. The pipeline for RB-TnSeq mutant fitness calculations is available at bitbucket.org/berkeleylab/feba/src/master. DIA-NN is freely available for download from https://github.com/vdemichev/DiaNN.

## Acknowledgements

We thank Chelsea Hernandez for assistance with exometabolomics sample processing and data collection, and Morgan Price for assistance with processing the RB-TnSeq data. This material is based upon work at the Joint BioEnergy Institute (JBEI) supported by the U.S. Department of Energy, Office of Science, Biological and Environmental Research Program under contract DE-AC02-05CH11231 with Lawrence Berkeley National Laboratory. This material by ENIGMA-Ecosystems and Networks Integrated with Genes and Molecular Assemblies (http://enigma.lbl.gov), a Science Focus Area Program at Lawrence Berkeley National Laboratory is based upon work supported by the U.S. Department of Energy, Office of Science, Office of Biological & Environmental Research under contract number DE-AC02-05CH11231. Metabolomics data analysis used resources of the National Energy Research Scientific Computing Center, a Department of Energy Office of Science User Facility. YW acknowledges the support from Texas A&M University and AgriLife Research.

## Author contributions

YW conceptualized the study and designed the experiments with input from TN. YW performed the primary experiments and initial data analysis. EP conducted additional experiments and led final data analysis with input from YW. EP and YW prepared the draft manuscript. YC and CP acquired proteomics data using LC-MS. VT assisted with the RB-TnSeq experiments, and VT and AD extracted DNA and acquired RB-TnSeq data. EP, MdR, and SK acquired metabolomics data using LC-MS/MS. RN performed flux balance analysis under the supervision of DS. TN, APA, and YW acquired funding. All authors revised the manuscript and approved the final version.

---

**Extended Data Figure 1.**
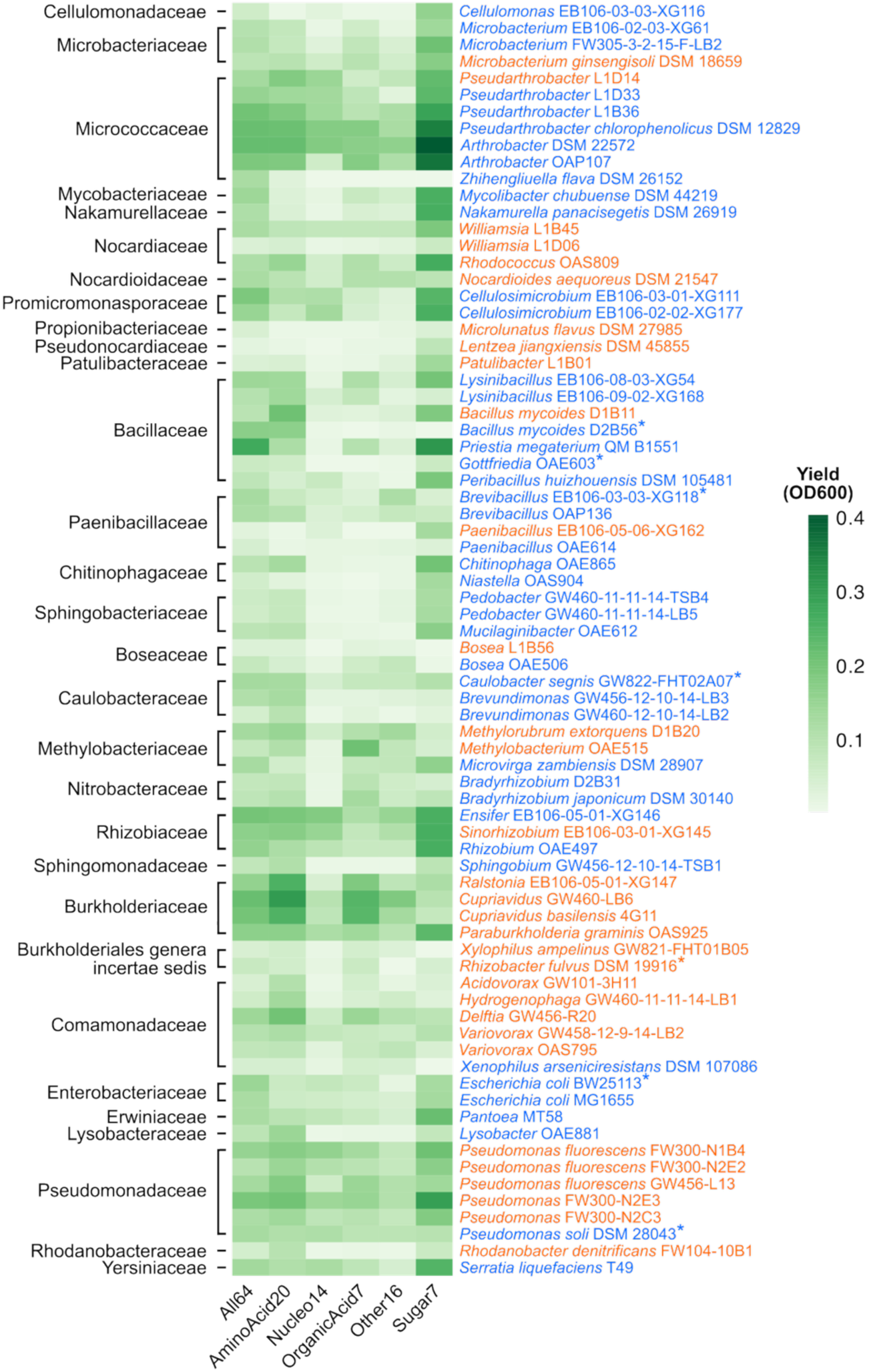
Growth yields of all tested strains on the experimental media. Heat map depicting the median growth yields of each tested strain in the six experimental media. Strains are grouped by taxonomic family, labeled on the left. The strain names are colored by the subsequently determined superadditive (blue) vs. additive (orange) growth phenotype in All64. Asterisks signify strains whose highest yield was in All64.

**Extended Data Figure 2.**
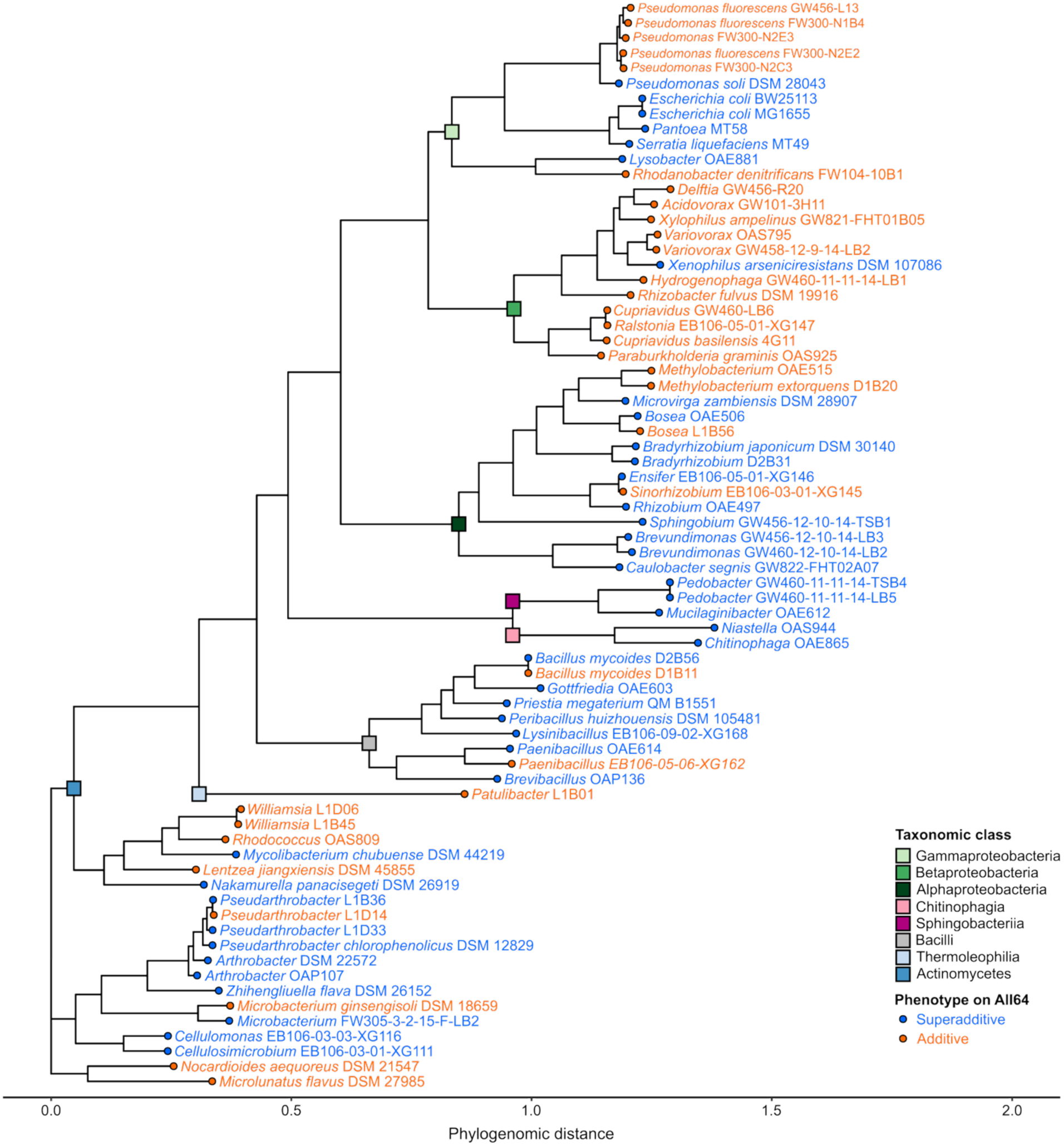
Phylogenomic tree of the 72 genome-sequenced isolates. The tree was constructed using default settings in GToTree based on the amino acid sequences of 74 conserved single-copy bacterial genes. The colored squares indicate taxonomic class (marked at the last common node for each class) and strain names and tip nodes are colored by growth phenotype in All64.

**Extended Data Figure 3.**
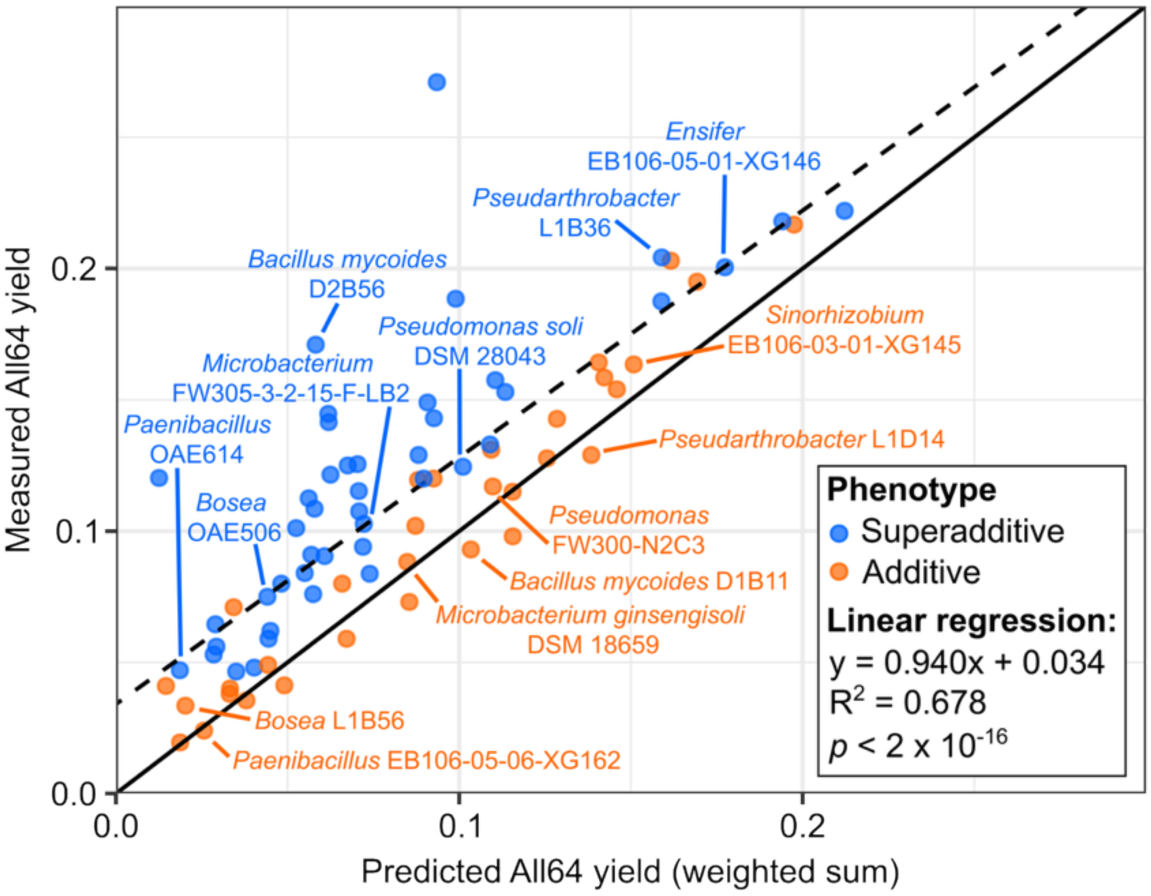
Measured versus predicted yield of closely-related strains with discordant growth phenotypes. Plot of measured growth yield on All64 versus the weighted-sum prediction as in Figure 1c, with labels highlighting pairs of closely related strains (same species or genus) that had contrasting phenotypes. Note that *Ensifer* and *Sinorhizobium* are synonymous genera. In cases where one strain within a genus was a contrast to all others in the genus, we have shown here only the two most closely related strains. The solid line is y = x and the dashed line is the best-fit linear regression model whose equation is given on the bottom right.

**Extended Data Figure 4.**
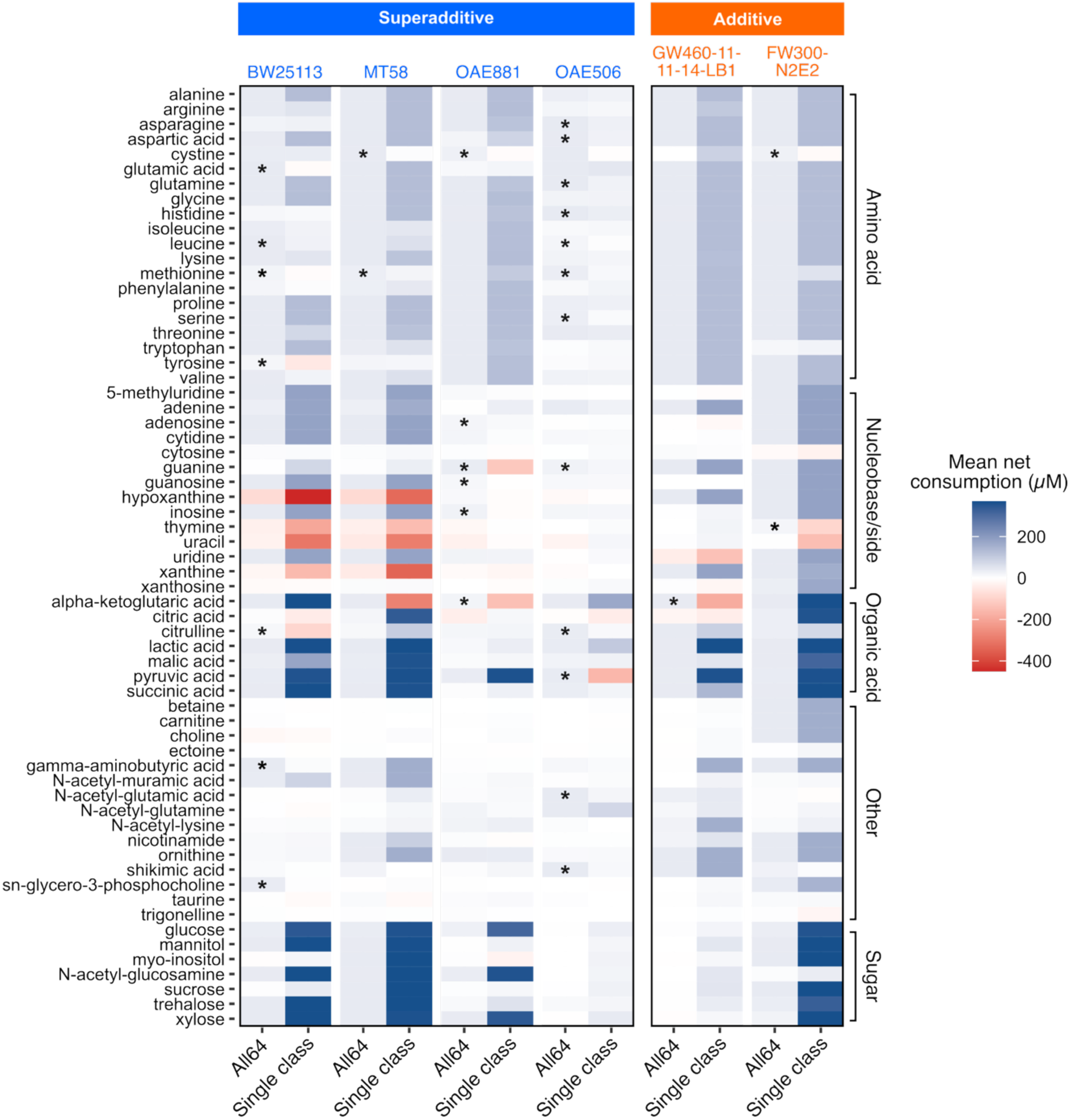
Net absolute depletion of supplied carbon sources from stationary phase supernatants. Heat map depicting the mean net consumption of 63 carbon sources in All64 versus the respective single-class medium (AminoAcid20, Nucleo14, OrganicAcid7, Other16, or Sugar7) in the supernatants of cultures grown to early stationary phase. Spermidine was excluded from the analysis as it was not detectable with the same chromatography method as all of the other carbon sources. Negative net consumption indicates net excretion, i.e. the concentration of the compound was higher in the supernatants than in the media controls. \**p* < 0.05 (Welch’s t-test followed by the Benjamini-Hochberg procedure), excluding compounds for which the percent remaining in All64 was either >100% or not significantly different from 100% (one-sample t-test).

**Extended Data Figure 5.**
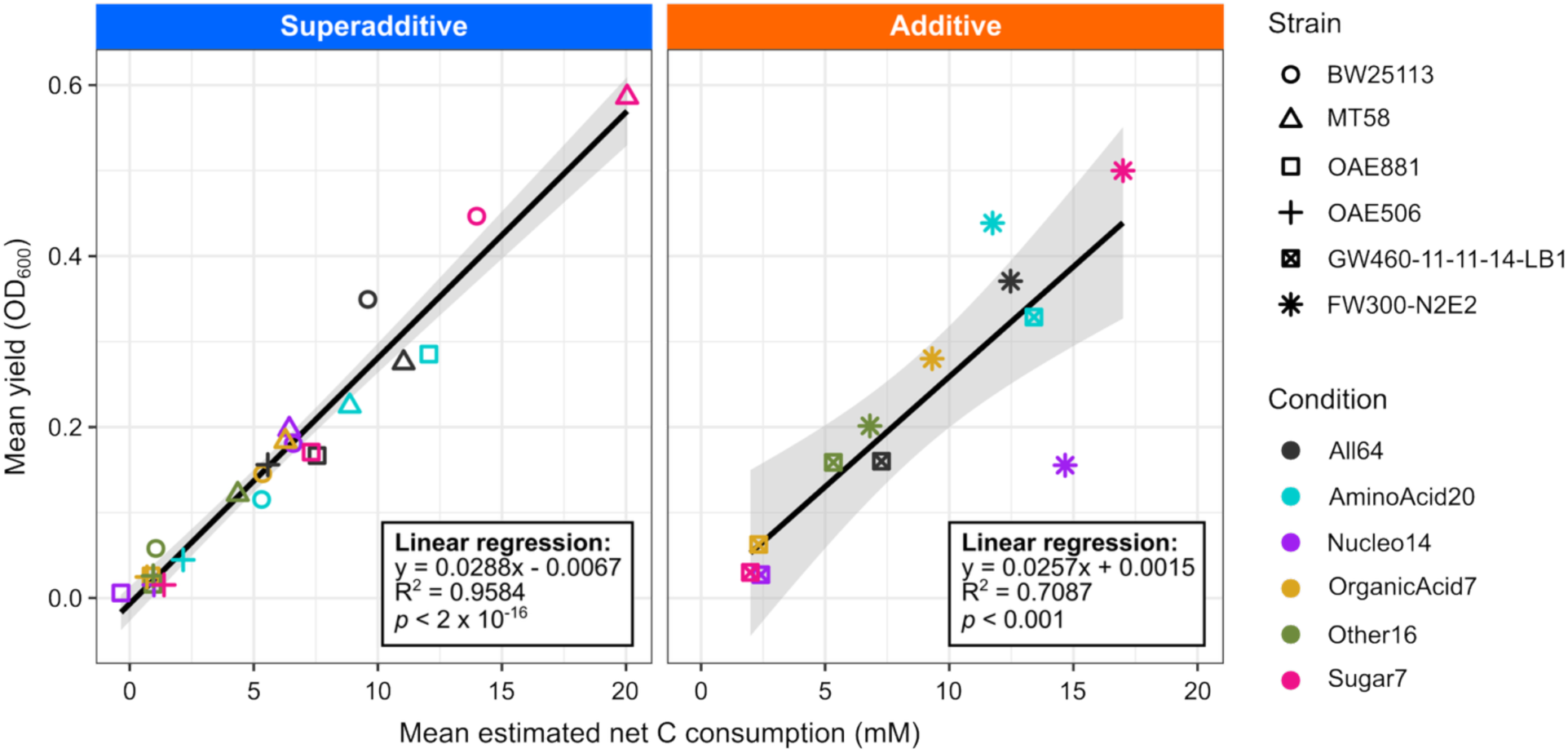
Growth yield correlates with estimated net carbon consumption. Growth yield (maximum OD_600_) of each strain plotted against the estimated total net amount of carbon (mM) depleted from the growth medium based on the net depletion of supplied compounds. Growth yields are from the same cultures as the supernatants used for exometabolomics, which were grown at a larger scale in flasks, rather than the original plate reader growth curves. The estimates for total net amount of carbon consumed do not account for any excreted organic compounds not originally present in the media. Each point represents the mean of three replicate cultures. The black lines show the best-fit linear regression for each group of strains, with the 95% confidence intervals shaded in gray. The slopes of the regression lines for superadditive versus additive growers are not significantly different (ANOVA *p*-value = 0.4582 for the interaction between growth phenotype and estimated net C consumption with growth yield as the response variable).

**Extended Data Figure 6.**
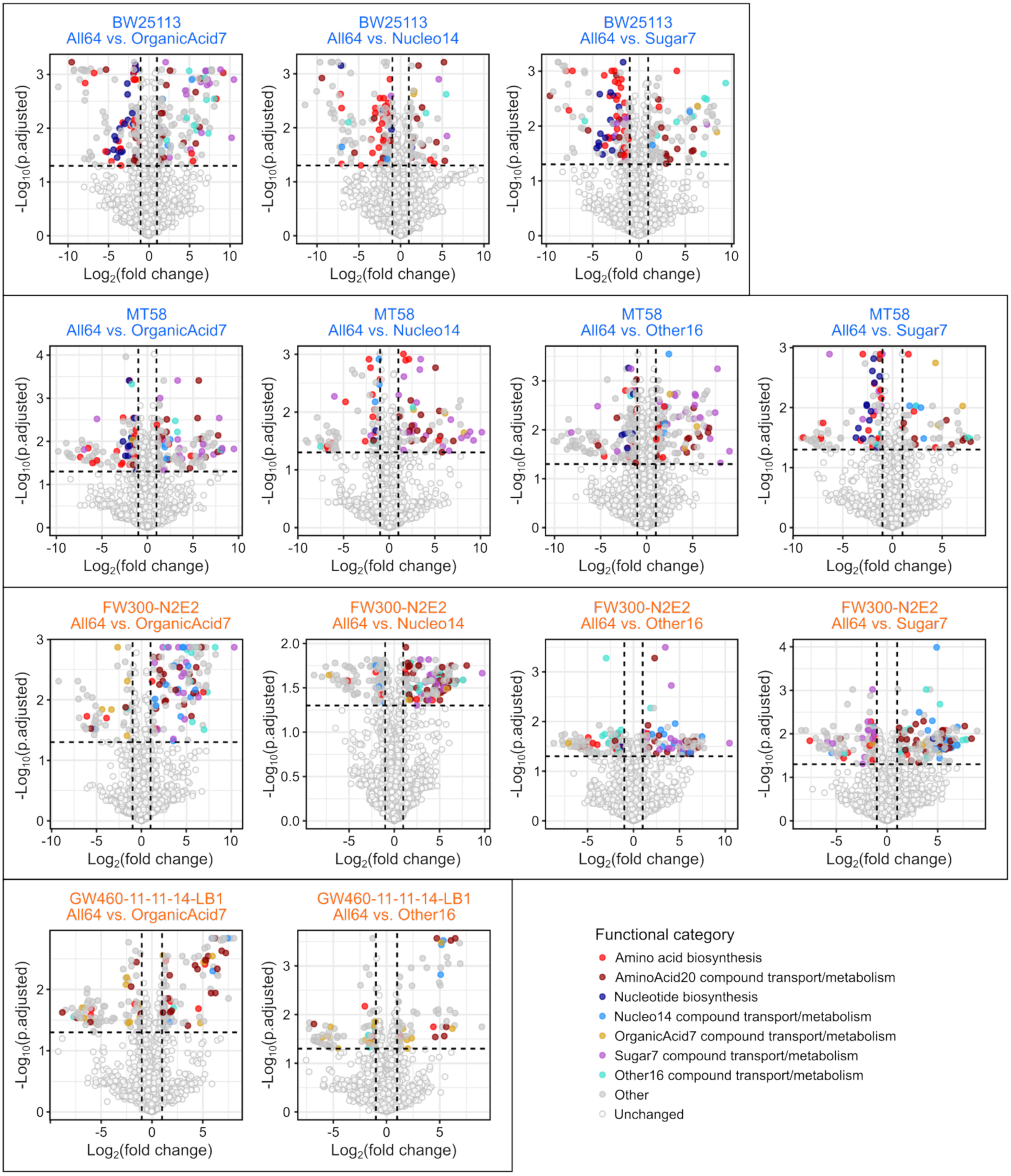
Differentially expressed proteins between All64 and single-class media for representative superadditive and additive strains. Proteomics volcano plots for All64 vs. OrganicAcid7, Nucleo14, Other16, and Sugar7 comparisons, organized by strain. *Bosea* OAE506 is not shown because it only produced enough biomass for proteomics in All64 and AminoAcid20, the results of which are depicted in Fig. 4b. Strain names are colored by phenotype (superadditive = blue, additive = orange).

**Extended Data Figure 7.**
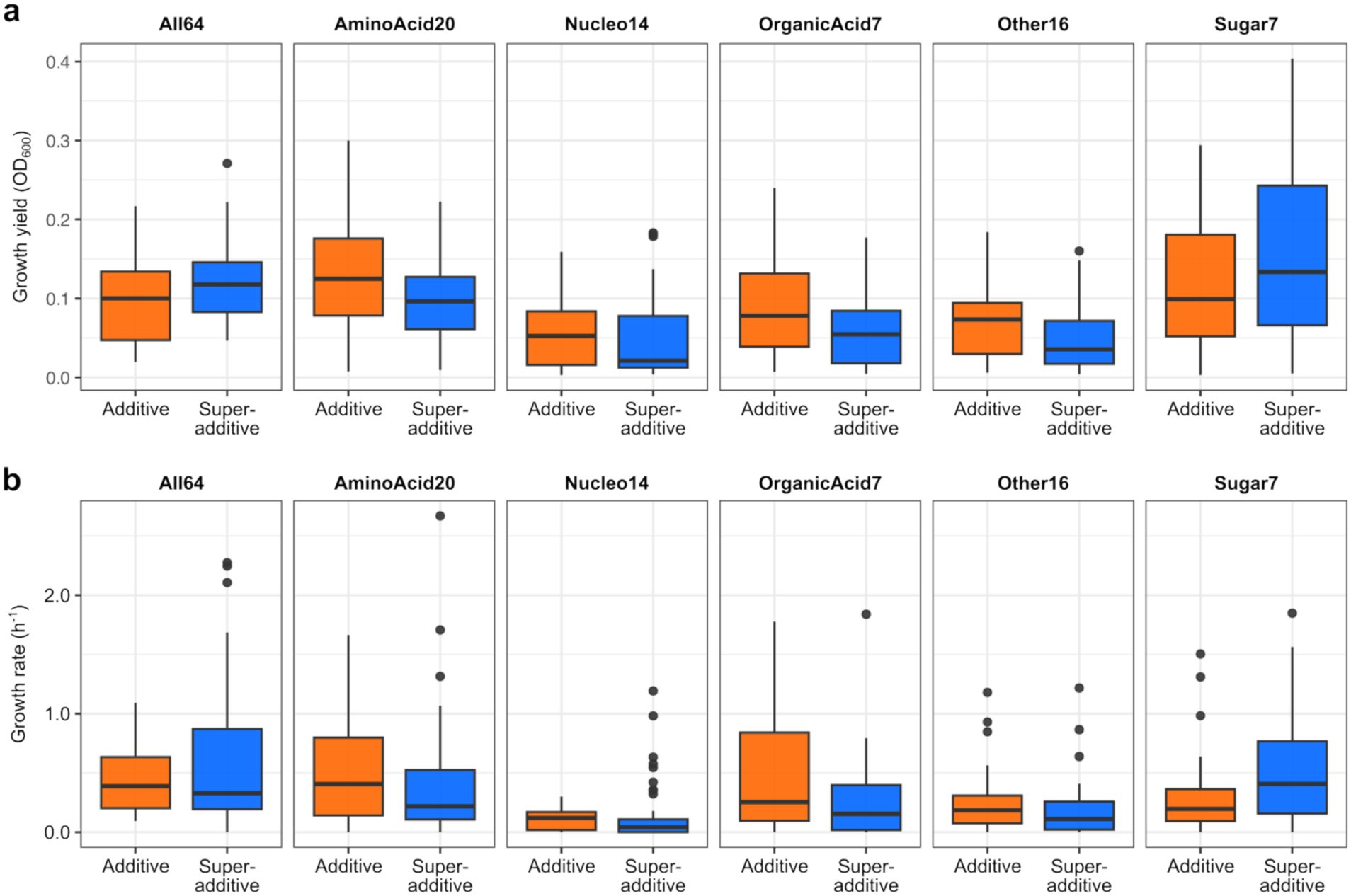
Growth yields and growth rates of superadditive versus additive growers across the experimental media. Growth yields (**a**) and growth rates (**b**) of all tested strains on the six experimental media, aggregated by growth phenotype on All64 (additive vs. superadditive). There were no statistically significant differences in growth yield or rate between the two phenotypic groups on any of the media (Mann-Whitney U test).

**Extended Data Figure 8.**
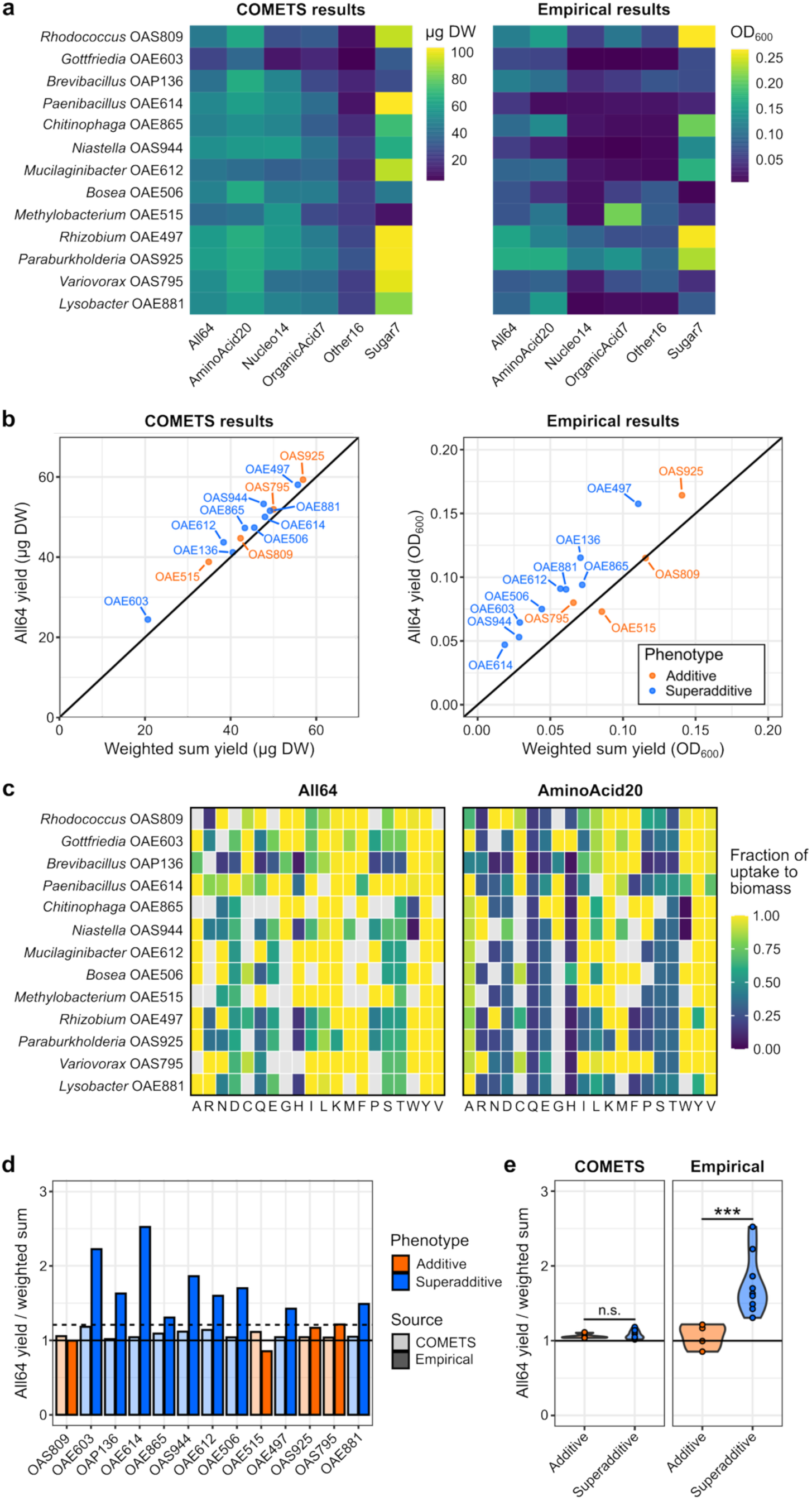
Flux-balance analysis with default parameters partially recapitulates superadditivity but fails to distinguish between superadditive and additive strains. **a)** Biomass yield predicted by COMETS in different media at 200 µL scale using genome-scale metabolic models for selected strains (left panel), compared against the empirical results (median of 3 replicates) for the same strains in the plate reader assay (right panel). Yield is given in units of µg dry weight (DW) for COMETS and OD600 for the empirical data. **b)** Plot of the All64 yield versus the weighted sum of the yields in the single-class media for the same strains as in (a), based on the COMETS simulations (left panel) or the empirical data (right panel). Data points in both panels are colored by the empirical phenotypic classification for each strain. **c)** Fraction of the uptake for each amino acid (labeled by single-letter codes on the horizontal axis) that COMETS predicted to be directed towards biomass for each strain at the first time-step in All64 vs. AminoAcid20. Blank tiles indicate that the amino acid was not predicted to be taken up or that the uptake reaction was absent from that GEM. **d)** Ratio of biomass produced on All64 versus the weighted sum prediction for the 14 strains with GEMs. Lighter colors represent the COMETS results and darker colors represent the empirical results. Orange (additive) and blue (superadditive) indicate the empirical classifications of the strains. The solid black line marks a ratio of 1 (i.e., exactly additive growth on All64). The dashed line marks the highest empirical ratio observed amongst the additive strains with GEMs, showing that none of the COMETS simulations exceeded this ratio, even for empirically superadditive strains. **e)** Violin plots showing that additive and superadditive growers (amongst strains with GEMs) were clearly separated by the empirical ratio of biomass produced on All64 versus the weighted sum prediction (*p* < 0.001, Welch’s t-test), whereas COMETS failed to recreate this difference.

## References

1. Monod, J. The growth of bacterial cultures. Annu. Rev. Microbiol. 3, 371–394 (1949).

2. Golan, O., Gampp, O., Eckert, L. & Sauer, U. Overall biomass yield on multiple nutrient sources. npj Syst. Biol. Appl. 11, 17 (2025).

3. Soong, J. L. et al. Microbial carbon limitation: the need for integrating microorganisms into our understanding of ecosystem carbon cycling. Glob. Change Biol. 26, 1953–1961 (2020).

4. Hobbie, J. E. & Hobbie, E. A. Microbes in nature are limited by carbon and energy: the starving-survival lifestyle in soil and consequences for estimating microbial rates. Front. Microbiol. 4, 324 (2013).

5. Bachmann, H., et al. Availability of public goods shapes the evolution of competing metabolic strategies. Proc. Natl. Acad. Sci. U.S.A. 110, 14302–14307 (2013).

6. Bickel, S. & Or, D. Aqueous habitats and carbon inputs shape the microscale geography and interaction ranges of soil bacteria. Commun. Biol. 6, 322 (2023).

7. Valiei, A., Dickson, A. M., Aminian-Dehkordi, J. & Mofrad, M. R. K. Bacterial community dynamics as a result of growth-yield trade-off and multispecies metabolic interactions toward understanding the gut biofilm niche. BMC Microbiol. 24, 441 (2024).

8. Liang, C., Schimel, J. P. & Jastrow, J. D. The importance of anabolism in microbial control over soil carbon storage. Nat. Microbiol. 2, 17105 (2017).

9. Jiao, N. et al. The microbial carbon pump and climate change. Nat. Rev. Microbiol. 22, 408–419 (2024).

10. Konzock, O. & Nielsen, J. TRYing to evaluate production costs in microbial biotechnology. Trends Biotechnol. 42, 1339–1347 (2024).

11. Pfleger, B. F. & Takors, R. Recent progress in the synthesis of advanced biofuel and bioproducts. Curr. Opin. Biotechnol. 80, 102913 (2023).

12. Fonte, E. S. et al. The combination of different carbon sources enhances bacterial growth efficiency in aquatic ecosystems. Microb. Ecol. 66, 871–878 (2013).

13. Fu, H., Uchimiya, M., Gore, J. & Moran, M. A. Ecological drivers of bacterial community assembly in synthetic phycospheres. Proc. Natl. Acad. Sci. U.S.A. 117, 3656–3662 (2020).

14. Pacheco, A. R., Moel, M. & Segrè, D. Costless metabolic secretions as drivers of interspecies interactions in microbial ecosystems. Nat. Commun. 10, 103 (2019).

15. de Raad, M. et al. A defined medium for cultivation and exometabolite profiling of soil bacteria. Front. Microbiol. 13, (2022).

16. Kell, D. B. et al. Metabolic footprinting and systems biology: the medium is the message. Nat. Rev. Microbiol. 3, 557–565 (2005).

17. Wetmore, K. M. et al. Rapid quantification of mutant fitness in diverse bacteria by sequencing randomly bar-coded transposons. mBio 6, e00306–15 (2015).

18. Price, M. N. et al. Mutant phenotypes for thousands of bacterial genes of unknown function. Nature 557, 503–509 (2018).

19. Thorgersen, M. P. et al. Deciphering microbial metal toxicity responses via random bar code transposon site sequencing and activity-based metabolomics. Appl. Environ. Microbiol. 87, e01037–21 (2021).

20. Price, M. N. et al. Novel enzymes for amino acid biosynthesis in diverse bacteria and archaea: GapMind 2026. 2024.10.14.618325 Preprint at 10.1101/2024.10.14.618325 (2026).

21. Kanehisa, M., Furumichi, M., Sato, Y., Matsuura, Y. & Ishiguro-Watanabe, M. KEGG: biological systems database as a model of the real world. Nucleic Acids Res. 53, D672–D677 (2025).

22. Li, Z., Selim, A. & Kuehn, S. Statistical prediction of microbial metabolic traits from genomes. PLoS Comput. Biol. 19, e1011705 (2023).

23. Gralka, M., Pollak, S. & Cordero, O. X. Genome content predicts the carbon catabolic preferences of heterotrophic bacteria. Nat. Microbiol. 8, 1799–1808 (2023).

24. Moyne, O. et al. Predicting competition and substrate preferences for targeted microbiome alteration. Cell 189, 3324–3338.e23 (2026).

25. Dukovski, I. et al. A metabolic modeling platform for the computation of microbial ecosystems in time and space (COMETS). Nat. Protoc. 16, 5030–5082 (2021).

26. Massucci, F. A., Guimerà, R., Nunes Amaral, L. A. & Sales-Pardo, M. Scaling and optimal synergy: Two principles determining microbial growth in complex media. Phys. Rev. E 91, 062703 (2015).

27. Fierer, N., Bradford, M. A. & Jackson, R. B. Toward an ecological classification of soil bacteria. Ecology 88, 1354–1364 (2007).

28. Malard, L. A. & Guisan, A. Into the microbial niche. Trends Ecol. Evol. 38, 936–945 (2023).

29. Pacheco, A. R., Osborne, M. L. & Segrè, D. Non-additive microbial community responses to environmental complexity. Nat. Commun. 12, 2365 (2021).

30. Shalev, O., Ye, X., Kilian, J., Stahl, M. & Ratzke, C. Increased nutrient diversity can induce loss of microbial diversity through enhanced resource uptake. Nat. Ecol. Evol. 10, 1449–1463 (2026).

31. Delgado-Baquerizo, M. et al. A global atlas of the dominant bacteria found in soil. Science 359, 320–325 (2018).

32. Baba, T. et al. Construction of Escherichia coli K-12 in-frame, single-gene knockout mutants: the Keio collection. Mol. Syst. Biol. 2, MSB4100050 (2006).

33. Wang, Y. et al. Substrate utilization and competitive interactions among soil bacteria vary with life-history strategies. Front. Microbiol. 13, (2022).

34. Reasoner, D. J. & Geldreich, E. E. A new medium for the enumeration and subculture of bacteria from potable water. Appl. Environ. Microbiol. 49, 1–7 (1985).

35. Blazanin, M. gcplyr: an R package for microbial growth curve data analysis. BMC Bioinform. 25, 232 (2024).

36. Sprouffske, K. & Wagner, A. Growthcurver: an R package for obtaining interpretable metrics from microbial growth curves. BMC Bioinform. 17, 172 (2016).

37. Lee, M. D. GToTree: a user-friendly workflow for phylogenomics. Bioinformatics 35, 4162–4164 (2019).

38. Yao, Y. et al. Analysis of metabolomics datasets with high-performance computing and metabolite atlases. Metabolites 5, 431–442 (2015).

39. Price, M. N., Deutschbauer, A. M. & Arkin, A. P. GapMind: automated annotation of amino acid biosynthesis. mSystems 5, 10.1128/msystems.00291-20 (2020).

40. Karp, P. D. et al. The BioCyc collection of microbial genomes and metabolic pathways. Brief. Bioinform. 20, 1085–1093 (2019).

41. Chen, Y., Gin, J. & Petzold, C. Alkaline-SDS cell lysis of microbes with acetone protein precipitation for proteomic sample preparation in 96-well plate format V.1. protocols.io 10.17504/protocols.io.6qpvr6xjpvmk/v1 (2023).

42. Chen, Y., Gin, J. & Petzold, C. Discovery proteomic (DIA) LC-MS/MS data acquisition and analysis V.2. protocols.io 10.17504/protocols.io.e6nvwk1z7vmk/v2 (2022).

43. Demichev, V., Messner, C. B., Vernardis, S. I., Lilley, K. S. & Ralser, M. DIA-NN: neural networks and interference correction enable deep proteome coverage in high throughput. Nat. Methods 17, 41–44 (2020).

44. Cantalapiedra, C. P., Hernández-Plaza, A., Letunic, I., Bork, P. & Huerta-Cepas, J. eggNOG-mapper v2: functional annotation, orthology assignments, and domain prediction at the metagenomic scale. Mol. Biol. Evol. 38, 5825–5829 (2021).

45. The Galaxy Community. Galaxy for accessible, reproducible, and collaborative data analyses: 2026 update. Nucleic Acids Res. 54, W105–W116 (2026).

46. Wright, M. N. & Ziegler, A. ranger: a fast implementation of random forests for high dimensional data in C++ and R. J. Stat. Softw. 77, 1–17 (2017).

47. Perez-Riverol, Y. et al. The PRIDE database resources in 2022: a hub for mass spectrometry-based proteomics evidences. Nucleic Acids Res. 50, D543–D552 (2022).

