## Supplementary Notes for "Diverse bacteria exhibit superadditive growth on mixed classes of carbon sources"

***1. Limitations of the exometabolomics analysis.***

An important caveat is that the disappearance of a compound from the medium does not necessarily mean that it was utilized as a carbon or energy source; it could be enzymatically transformed or taken up into the cells without being further metabolized, which our assay would not reveal. Additionally, for many compounds, both consumption and excretion could potentially have contributed to the observed final concentration. We also did not attempt to detect or quantify any compounds that were not originally provided in the media but may have been excreted as metabolic by-products. Despite these methodological limitations, we still observed strong correlations between total estimated net carbon consumption and growth yield, with similar slopes for both additive and superadditive growers. The only clear outlier was the relatively low yield of *P. fluorescens* FW300-N2E2 on Nucleo14. However, those samples were collected at the first plateau of a diauxic growth curve rather than the final maximum OD600, unlike all other samples, so this lower-than-expected yield may simply reflect as-yet-incomplete processing of the depleted nucleobases/sides into biomass.

***2. Fitness effects of genes related to carbon source catabolism.***

A neutral fitness score for a catabolic gene does not necessarily indicate that the respective carbon source was not used, given that in a medium like All64 where many carbon sources are available, loss of the ability to consume any one in particular might not significantly impact overall fitness. Nevertheless, for three of the four superadditive growers (*E. coli* BW25113, *Pantoea* MT58, and *Lysobacter* OAE881), multiple genes related to the transport and/or degradation of sugars and amino acids met our criteria for significance ($|t| > 4$ and fitness score $< -1$), suggesting that catabolism of both classes of substrates was important for full fitness of these strains in All64. The fourth superadditive grower, *Bosea* OAE506, also had significant negative fitness scores in All64 for multiple genes related to amino acid degradation, but unlike in the other superadditive strains, no genes definitively related to sugar catabolism had negative fitness scores. This is not surprising as *Bosea* OAE506 failed to grow on Sugar7 and exhibited little to no consumption of sugars in the exometabolomics experiment (Fig. 2).

For the additive growers, only a few genes clearly related to carbon source catabolism had significant negative fitness scores in All64. In *Hydrogenophaga* GW460-11-11-14-LB1, methylmalonyl-CoA mutase and glucose 6-phosphate isomerase were the only two such genes. Methylmalonyl-CoA mutase participates in degradation of branched-chain amino acids, suggesting that these may have been a particularly important carbon source for this strain. Glucose 6-phosphate isomerase participates in both glycolysis and gluconeogenesis but was most likely playing the latter role in this strain, since it consumed amino acids and organic acids to a much greater extent than sugars in the exometabolomics experiment (Fig. 2). In *P. fluorescens* FW300-N2E2, on the other hand, the three genes with significant negative fitness scores in All64—sucrose 6-phosphate hydrolase, glucose-6-phosphate 1-dehydrogenase, and fumarylacetoacetase (involved in phenylalanine and tyrosine degradation)—suggested that similar to some of the consistently superadditive growers, this strain relied on catabolism of both sugars and amino acids in All64.

***3. Evidence for regulatory inefficiencies based on RB-TnSeq.***

Regulatory inefficiencies in the additive strains are suggested by the observation of genes with significant positive fitness scores specific to All64—that is, genes with $|t| > 4$ and a fitness score $> 1$ in All64 and $< 0.5$ in all other media. As noted in the main text, these genes were pyruvate dehydrogenase E1 component in *Hydrogenophaga* GW460-11-11-14-LB1, and a maltose/maltodextrin transporter in *P. fluorescens* FW300-N2E2. Pyruvate dehydrogenase was actually required for the fitness of *Hydrogenophaga* GW460-11-11-14-LB1 (i.e. had a strong negative fitness score) in most of the other media except for AminoAcid20, where its effect was neutral, consistent with its known role as a central gene connecting glycolysis and the TCA cycle. Yet its detrimental effect in All64 was extremely strong (fitness score of 6.5 with $|t| > 37$), suggesting that *Hydrogenophaga* GW460-11-11-14-LB1 may have expressed this gene at a high level, and consequently high cost, in All64 despite apparently not benefitting from its function under this condition. Similarly, the detrimental effect of the maltose/maltodextrin transporter in *P. fluorescens* FW300-N2E2 may suggest that this gene was inappropriately induced by other sugars present in All64, which does not contain maltose, thus costing the cells metabolically without providing a benefit. Besides these genes, the nucleotide biosynthesis genes pyrB, pyrC, and pyrD in *Hydrogenophaga* GW460-11-11-14-LB1 also met our fitness score cutoffs for a specific positive fitness score in All64. While the $|t|$ scores for these genes were below our cutoff for statistical significance, this result hints that *Hydrogenophaga* GW460-11-11-14-LB1 is able to salvage pyrimidine nucleobases but fails to downregulate the respective de novo biosynthesis genes accordingly (the latter was subsequently confirmed by proteomics), thus incurring a fitness cost when exogenous nucleobases are available. Ultimately, however, a positive fitness effect in a pooled assay does not identify whether the relevant cost is protein expression, altered flux, a toxic intermediate, or a polar effect.
