## Supplementary Discussion for "Diverse bacteria exhibit superadditive growth on mixed classes of carbon sources"

### *1. Relative importance of different putative mechanisms for superadditive growth.*

What is the relative importance of the putative physiological mechanisms we have identified for superadditive growth? We speculate that increased carbon uptake when different classes of carbon sources are mixed is more important overall for superadditive growth than downregulation of biosynthetic genes, based on a few observations: 1) the relationship between estimated total net C consumption and growth yield was remarkably tight across different strains and media in the exometabolomics experiment, with a plausible technical explanation for the only obvious outlier; 2) the proteomics results for *Bosea* sp. OAE506 suggest that downregulation of biosynthetic pathways may not be a universal trait of superadditive growers (though we could not assess the regulation of amino acid biosynthesis in this strain); and 3) the main repressor of purine biosynthesis in Enterobacterales members is well-known, but this gene (*purR*) did not have a significant fitness effect in All64 for either *E. coli* BW25113 or *Pantoea* MT58. However, with regard to the latter point, it is possible that losing the downregulation of just one biosynthetic pathway amongst several downregulated pathways is insufficient to cause a serious fitness defect in All64.

Indeed, while increased carbon uptake may be more broadly linked to superadditive growth across diverse strains, our results nevertheless suggest that downregulation of biosynthetic enzymes also contributes to the magnitude of extra biomass gain during growth on mixed classes of carbon sources, and that different combinations of these mechanisms may lead to similar results. For example, while *Bosea* sp. OAE506 did not downregulate nucleotide biosynthesis in All64 relative to AminoAcid20, it consumed more carbon than expected in All64 by one of the largest absolute margins and by far the largest percentage difference among the six strains used for exometabolomics. Meanwhile, *Pantoea* sp. MT58 and *Lysobacter* sp. OAE881 exhibited both a moderate increase in consumed carbon in All64 and a moderate number of dependencies on biosynthetic genes/pathways in the RB-TnSeq experiments (i.e., more than *E. coli* BW25113 but less than *Bosea* sp. OAE506). All three of these strains (*Bosea* sp. OAE506, *Pantoea* sp. MT58, *Lysobacter* sp. OAE881) fall on the regression line in Fig. 1c, indicating that they all exceeded the weighted sum biomass prediction by an average amount even though their strategies apparently differ. *E. coli* BW25113, on the other hand, consumed additional carbon in All64 by one of the largest absolute margins, had the fewest dependencies on biosynthetic genes/pathways, and downregulated more biosynthetic genes in All64 than *Pantoea* MT58 or *Bosea* OAE506, and it correspondingly falls well above the regression line. While our representative strains did not happen to include one that downregulates numerous biosynthetic pathways in All64 while only consuming approximately the expected amount of carbon, we speculate that such a strain would also exhibit superadditive growth, but not to the extent of *E. coli* BW25113.

Finally, there are also other potential mechanisms that could contribute to superadditive growth that we have not addressed in this study. For example, during growth on mixed substrates, some substrates might be used as an energy source to enhance assimilation of carbon from other substrates, thereby boosting overall biomass yield[48]. Investigating this possibility would require the use of stable-isotope probing to track the fates of different substrates in single- versus mixed-class media, which could be a worthwhile endeavor in the future.

***2. Limitations of the definition of superadditive growth in this study.***

It is important to be clear about what superadditive growth does not necessarily imply. Our definition of superadditive growth strictly refers to whether a strain's growth yield on a diverse mixture of carbon sources exceeds the weighted sum of its growth on subsets of those carbon sources. Because the definition is based on a within-strain comparison, classification of one strain as a superadditive grower and another as an additive grower does not indicate which one will achieve a higher yield in a given condition. Indeed, while we found a slight trend towards superadditive growers achieving higher yields on All64 than additive growers, the overall difference is not statistically significant (Extended Data Fig. 7a). Similarly, there was no significant difference between superadditive and additive growers in terms of growth rate on All64, albeit the superadditive group had a longer tail of fast growers (Extended Data Fig. 7b). Furthermore, most superadditive growers achieved their highest growth yields on a medium other than All64 (Extended Data Fig. 1), so it is not generally true that a diverse but dilute mixture of carbon sources improves the biomass yield of a superadditive grower over a more-concentrated mixture of their preferred type of carbon source. Furthermore, we did not measure respiration or the total carbon content of the biomass produced, so it remains unclear whether the carbon use efficiency of superadditive growers differs from additive growers on mixtures of diverse carbon sources, although the consistent correlation we observed between estimated total net carbon consumption and optical density, regardless of phenotype (Extended Data Fig. 5), suggests that a systematic difference may be unlikely.

***3. Other potential uses of energy savings conferred by downregulation of biosynthetic pathways.***

It is well-established that loss or downregulation of biosynthetic pathways confers energy and proteomic resource savings and can therefore be adaptive when the products of those pathways are available externally[49]; indeed, such benefits are thought to frequently drive reductive genome evolution in free-living bacteria, as described by the “Black Queen

Hypothesis"[50]. However, the consistent relationship between estimated net carbon consumption and optical density across different strains and media suggests that not all of the biosynthetic cost savings facultatively realized by some superadditive growers are routed into increased biomass production per unit of consumed carbon. This raises the question of how else those resources may be spent. Possibilities for this include increased motility or increased production of secondary metabolites, both of which could in turn affect interspecies interactions and community structure. In support of the first possibility, All64 upregulated the expression of multiple flagellar components and chemotaxis proteins in the superadditive growers *E. coli* BW25113 and *Pantoea* MT58 compared to most single-class media (Supplementary Table 7), whereas this was generally not true (or even the opposite) for the two representative additive growers, nor for *Bosea* OAE506, which is a superadditive grower but for which we do not have evidence that downregulation of biosynthetic pathways contributes to its growth phenotype.

***4. Limitations of the study design for mechanistic inference and key open questions.***

Since the concentration of each metabolite in All64 is lower versus the single-class media we cannot distinguish whether the greater-than-expected uptake of certain substrates in All64 was driven by the presence of substrates from other classes, by the differences in concentration (e.g. non-linear scaling at low concentrations), or both. Since the same limitation applies to other recent multi-strain studies on mixed substrate growth[29,30], testing these factors separately should be a high priority for future work. Meanwhile, our observations regarding the differential importance and expression of biosynthetic pathways in All64 across different strains, regardless of their ability to take up the respective metabolites, suggest that there is still much to be learned about the regulation of amino acid and nucleobase/side biosynthesis in diverse taxa. Despite an ever-growing body of literature on carbon catabolite repression[51,52], there remain many knowledge gaps with regard to how, when, and why different species choose to route such compounds into catabolic versus anabolic pathways, especially outside of a few relatively well-studied model organisms. Further complicating matters is that different molecular mechanisms can lead to similar results, as in the case of the regulation of *de novo* purine biosynthesis in *E. coli* versus *Bacillus subtilis*[53,54]. Such heterogeneity in molecular mechanisms, not all of which are captured by protein sequences, may account in part for the limitations of our random forest classifier trained on KEGG ortholog presence/absence.

35. Blazanin, M. gcplyr: an R package for microbial growth curve data analysis. *BMC Bioinform.* **25**, 232 (2024).
36. Sprouffske, K. & Wagner, A. Growthcurver: an R package for obtaining interpretable metrics from microbial growth curves. *BMC Bioinform.* **17**, 172 (2016).
37. Lee, M. D. GToTree: a user-friendly workflow for phylogenomics. *Bioinformatics* **35**, 4162–4164 (2019).
38. Yao, Y. *et al.* Analysis of metabolomics datasets with high-performance computing and metabolite atlases. *Metabolites* **5**, 431–442 (2015).
39. Price, M. N., Deutschbauer, A. M. & Arkin, A. P. GapMind: automated annotation of amino acid biosynthesis. *mSystems* **5**, 10.1128/msystems.00291-20 (2020).
40. Karp, P. D. *et al.* The BioCyc collection of microbial genomes and metabolic pathways. *Brief. Bioinform.* **20**, 1085–1093 (2019).
41. Chen, Y., Gin, J. & Petzold, C. Alkaline-SDS cell lysis of microbes with acetone protein precipitation for proteomic sample preparation in 96-well plate format V.1. *protocols.io* https://dx.doi.org/10.17504/protocols.io.6qpvr6xjpvmk/v1 (2023).
42. Chen, Y., Gin, J. & Petzold, C. Discovery proteomic (DIA) LC-MS/MS data acquisition and analysis V.2. *protocols.io* https://dx.doi.org/10.17504/protocols.io.e6nvwk1z7vmk/v2 (2022).
43. Demichev, V., Messner, C. B., Vernardis, S. I., Lilley, K. S. & Ralser, M. DIA-NN: neural networks and interference correction enable deep proteome coverage in high throughput. *Nat. Methods* **17**, 41–44 (2020).
44. Cantalapiedra, C. P., Hernández-Plaza, A., Letunic, I., Bork, P. & Huerta-Cepas, J. eggNOG-mapper v2: functional annotation, orthology assignments, and domain prediction at the metagenomic scale. *Mol. Biol. Evol.* **38**, 5825–5829 (2021).
45. The Galaxy Community. Galaxy for accessible, reproducible, and collaborative data analyses: 2026 update. *Nucleic Acids Res.* **54**, W105–W116 (2026).
46. Wright, M. N. & Ziegler, A. ranger: a fast implementation of random forests for high dimensional data in C++ and R. *J. Stat. Softw.* **77**, 1–17 (2017).
47. Perez-Riverol, Y. *et al.* The PRIDE database resources in 2022: a hub for mass spectrometry-based proteomics evidences. *Nucleic Acids Res.* **50**, D543–D552 (2022).
48. Harder, W. & Dijkhuizen, L. Strategies of mixed substrate utilization in microorganisms. *Philos. Trans. R. Soc. Lond. B Biol. Sci.* **297**, 459–480 (1982).
49. D'Souza, G. *et al.* Less is more: selective advantages can explain the prevalent loss of biosynthetic genes in bacteria. *Evolution* **68**, 2559–2570 (2014).
50. Morris, J. J., Lenski, R. E. & Zinser, E. R. The Black Queen Hypothesis: evolution of dependencies through adaptive gene loss. *mBio* **3**, 10.1128/mbio.00036-12 (2012).
51. Görke, B. & Stülke, J. Carbon catabolite repression in bacteria: many ways to make the most out of nutrients. *Nat. Rev. Microbiol.* **6**, 613–624 (2008).
52. Park, H., McGill, S. L., Arnold, A. D. & Carlson, R. P. Pseudomonad reverse carbon catabolite repression, interspecies metabolite exchange, and consortial division of labor. *Cell. Mol. Life Sci.* **77**, 395–413 (2019).
53. Ayoub, N., Gedeon, A. & Munier-Lehmann, H. A journey into the regulatory secrets of the de novo purine nucleotide biosynthesis. *Front. Pharmacol.* **15**, (2024).
54. Sinha, S. C. *et al.* The purine repressor of *Bacillus subtilis*: a novel combination of domains adapted for transcription regulation. *J. Bacteriol.* **185**, 4087–4098 (2003).
